# Systematic discovery of gene fusions in pediatric cancer by integrating RNA-seq and WGS

**DOI:** 10.1101/2021.08.31.458342

**Authors:** Ianthe A.E.M. van Belzen, Casey Cai, Marc van Tuil, Shashi Badloe, Eric Strengman, Alex Janse, Eugène T. Verwiel, Douwe F.M. van der Leest, Lennart Kester, Jan J. Molenaar, Jules Meijerink, Jarno Drost, Weng Chuan Peng, Hinri H.D. Kerstens, Bastiaan B.J. Tops, Frank C.P. Holstege, Patrick Kemmeren, Jayne Y. Hehir-Kwa

**Affiliations:** Princess Máxima Center for Pediatric Oncology, Utrecht, The Netherlands; Department of Pharmaceutical Sciences, Utrecht University, The Netherlands; Oncode Institute, Utrecht, The Netherlands

**Keywords:** Gene fusions, chimeric transcripts, pediatric cancer, structural variants, whole genome sequencing, rna sequencing

## Abstract

**Background:** Gene fusions are important cancer drivers in pediatric cancer and their accurate detection is essential for diagnosis and treatment. Clinical decision-making requires high confidence and precision of detection. Recent developments show RNA sequencing (RNA-seq) is promising for genome-wide detection of fusion products, but hindered by many false positives that require extensive manual curation and impede discovery of pathogenic fusions.

**Results:** We developed Fusion-sq to detect tumor-specific gene fusions by integrating and “fusing” evidence from RNA-seq and whole genome sequencing (WGS) using intron-exon gene structure. In a pediatric pan-cancer cohort of 130 patients, we identified 165 high confidence tumor-specific gene fusions and their underlying structural variants (SVs). This includes all clinically relevant fusions known to be present in this cohort (30 patients). Fusion-sq distinguishes healthy-occurring from tumor-specific fusions, and resolves fusions in amplified regions and copy number unstable genomes. A high gene fusion burden is associated with copy number instability. We identified 27 potentially pathogenic fusions involving oncogenes or tumor-suppressor genes characterised by underlying SVs or expression changes indicative of activating or disruptive effects.

**Conclusions:** Our results indicate how clinically relevant and potentially pathogenic gene fusions can be identified and their functional effects investigated by combining WGS and RNA-seq. Integrating RNA fusion predictions with underlying SVs advances fusion detection beyond extensive manual filtering. Taken together, we developed a method for identifying candidate fusions that is suitable for precision oncology applications. Our method provides multi-omics evidence for assessing the pathogenicity of tumor-specific fusions for future clinical decision making.

## Background

Gene fusions are important driver mutations in cancer and their accurate detection is essential for diagnosis, treatment selection and understanding disease mechanisms. The fusion of two or more genes through a structural variant (SV) can affect the involved genes directly, but also give rise to a chimeric protein with oncogenic properties [1, 2]. SVs can dysregulate cells in multiple ways, for instance by disrupting genes or by displacing an enhancer resulting in overexpression of oncogenes (e.g. *TLX1/3, NKX2-1*)[3]. However, gene fusions are a distinct type of variants characterised by the formation of fusion products and their chimeric transcripts[3]. The contribution of gene fusions to cancer etiology strongly differs per cancer type and they are especially important in the diagnostic process of pediatric cancers [1, 2]. For example, *KIAA1549--BRAF* fusions in pilocytic astrocytoma and *EWRS1--FLI* fusions in Ewing sarcoma are prime determinants of these tumor types[2]. Also many leukemias have characteristic driver fusions [4]. Detection of these fusions for diagnostic purposes is usually done with targeted assays, which are reliable, fast and cost-effective, but limited to known partner genes and/or breakpoints[5, 6]. As a result, targeted assays fail to detect some fusions with alternative breakpoints (e.g. *KIAA1549--BRAF)* [7] *or which have different partner genes (e*.*g. fusions involving TFE3, NUP98, FGFR*)[8–10]. These limitations also make targeted assays unsuitable for discovery of novel gene fusions.

RNA sequencing (RNA-seq) is increasingly applied in research to detect gene fusions that result in chimeric transcripts. More recently within a diagnostic context, it has been shown that RNA-seq is a robust alternative to targeted assays for detecting “clinically relevant” gene fusions. Here we consider fusions as clinically relevant if they have been published in peer-reviewed journals as associated with specific cancer types and are used for diagnosis, prognosis and treatment selection. In a pediatric cancer cohort, a 38% increase in diagnostic yield was achieved with RNA-seq compared to traditional diagnostic assays (Hehir-kwa, 2021, manuscript under revision). However, one of the key issues with robust detection of gene fusions based solely on RNA-seq data is controlling the false positive rate. Hundreds of chimeric transcripts per sample can be detected by RNA-seq and predicted to reflect gene fusions [11]. Although some have underlying genomic SVs, others result from normal transcription processes such as read-through and intergenic trans-splicing events [12]. Many chimeric transcripts are found in healthy tissue with no known links to malignancy [12, 13]. For example, the translocation resulting in a *PAX3--FOXO1* gene fusion is a driver mutation in alveolar rhabdomyosarcoma, but the chimeric transcript is also transiently expressed in healthy muscle tissue without an underlying SV[12, 13]. Therefore, fusion predictions from RNA-seq require stringent filtering to remove technical artefacts and healthy-occurring chimeric transcripts, as well as to rescue detection of lowly expressed fusions known to be clinically relevant. This introduces bias in the detection as the filtering is done with manually curated inclusion and exclusion lists which also limits its use for gene fusion discovery purposes. Alternatively, identifying the underlying SV can distinguish bonafide tumor-specific gene fusions from artefacts and other chimeric transcripts, as well as provide support to lowly expressed chimeric transcripts without the need for biased manual filtering.

Combining RNA-seq with SVs inferred from whole genome sequencing (WGS) data can help to detect potentially pathogenic gene fusions by identifying breakpoints that support the genomic origin of chimeric transcripts [14]. These SVs can be classified as tumor-specific based on analysis of paired tumor and normal WGS samples, whilst matching normal tissue for RNA isolation is often problematic to obtain. By itself, WGS is less suitable for reliable detection of actively transcribed gene fusions as SVs can affect multiple genes and WGS also infers many other non-transcribed variants. Similar to RNA-seq, WGS is prone to technical artifacts and false positives [15]. Despite this, using these orthogonal sequencing methods in combination is promising for genome-wide gene fusion detection as demonstrated for large cancer cohorts [14, 16].

To identify and interpret tumor-specific gene fusions in individual patients, we resolved the underlying SVs by combined analysis of RNA-seq and WGS data. For a heterogeneous cohort of 130 pediatric cancer patients, SVs were matched to gene fusion predictions based on intron-exon gene structure. For all 30 patients in which clinically relevant fusions were detected by RNA-seq, the underlying tumor-specific SVs were resolved with high confidence. Our approach avoids filtering out lowly expressed fusions while still removing healthy-occurring chimera. In 34 other patients without a known clinically relevant fusion, we detected 126 tumor-specific fusions with similar high confidence, including 27 distinct fusions involving oncogenes or tumor-suppressor genes. To further assess their potential pathogenicity, we analyzed the characteristics of these underlying SVs and changes in gene expression of the fusion partner genes. We found an association between copy number gain and overexpression, and cases of potential oncogene activation or tumor-suppressor gene disruption. In conclusion, we show that integration of RNA-seq and WGS data can be used to identify tumor-specific fusions. Also, the multi-omics evidence gathered by Fusion-sq about these gene fusions can aid molecular tumor boards in assessing their potential pathogenicity.

## Results

### Fusion-sq: detecting tumor-specific gene fusions with high confidence

To resolve gene fusions and investigate their relevance to pediatric cancer, we combined RNA-seq and paired tumor and normal WGS samples of 130 patients across 53 pediatric cancer types (Fig. 1a). Known clinically relevant fusions were identified with RNA-seq in 30 patients (Fig. 1a, grey overlay). We investigated the possibility of combining WGS structural variant analysis with RNA-seq data to increase detection specificity and identify potentially pathogenic gene fusions in the remainder of the cohort. Hereto, we developed Fusion-sq which integrates (“fuses”) predicted gene fusions from RNA-seq data with SVs from WGS data (Methods, Fig. 1b,c). For every predicted gene fusion, Fusion-sq first derives genomic intervals to match RNA and DNA breakpoints based on intron-exon gene structure. Next, DNA breakpoints falling within these intervals are used to identify SVs that link the fusion 5’ and 3’ partner genes. To optimize both recall and precision, SVs detected by Manta [17], DELLY [18] and GRIDSS [19] are considered separately and their support is summarized per fusion. We further selected high confidence fusions supported by SVs that are detected by at least two tools and correspond to the chimeric transcript (Fig. 1b,d, Additional file 1: Figure S1). Fusions are then classified as tumor-specific, (likely) germline or low allele fraction based on the SV’s variant supporting reads in the paired tumor and normal samples, as reflected in the allele fractions (AF) of SVs in both samples (Fig. 1b,d). In total, 16,144 fusions were predicted in 130 patients (median 101) using RNA-seq data alone (Fig. 1b). By combining these fusion predictions with WGS data, Fusion-sq identified 359 fusions supported by at least one SV tool, further refined into 232 high confidence fusions in 86 patients (median 1)(Fig. 1b,e). To investigate potentially pathogenic gene fusions, we further analyzed the 165 high confidence tumor-specific fusions (hcTSF) by combining RNA evidence with properties of the underlying SVs, such as SV type and allele fraction.

**Fig. 1:**
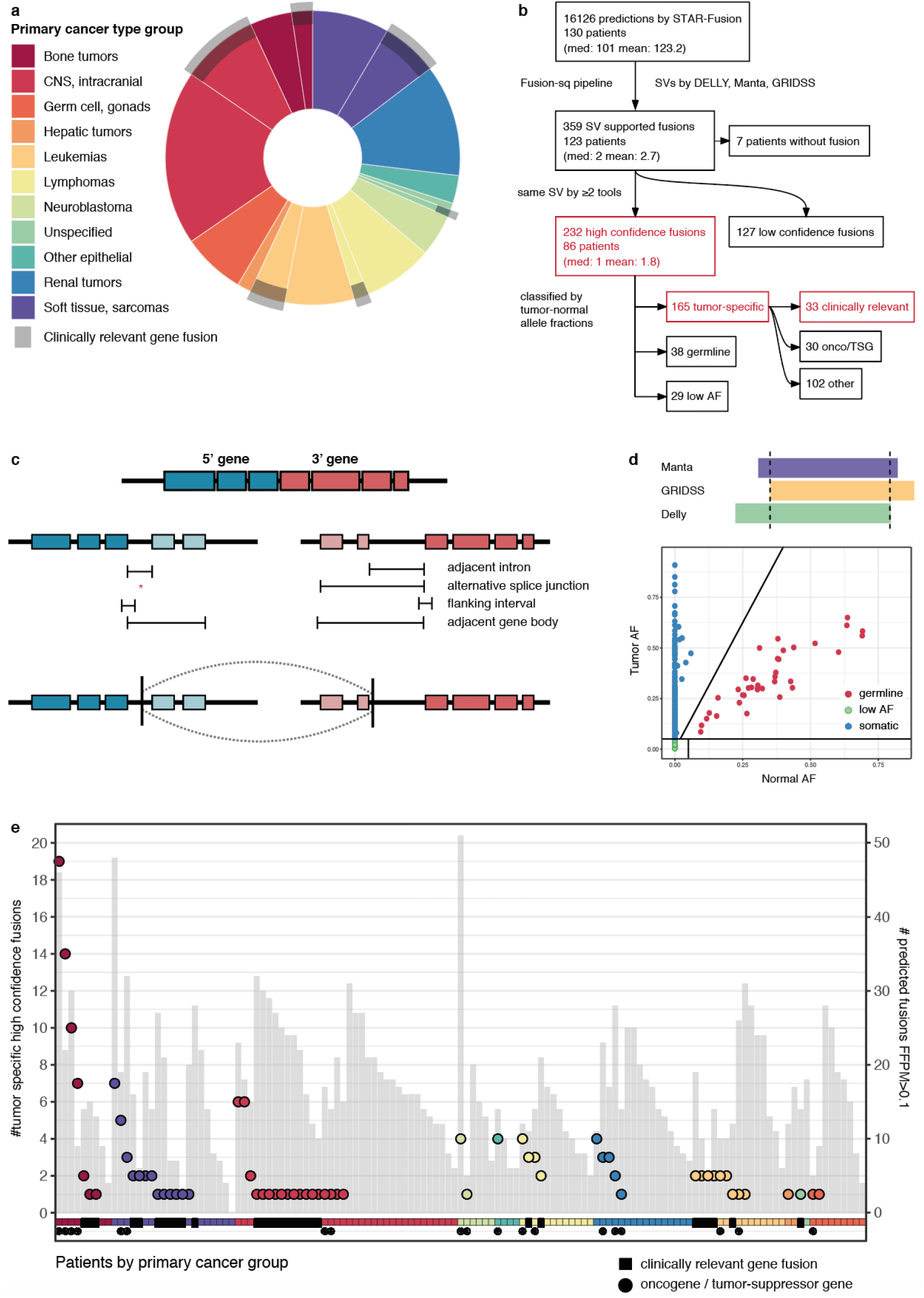
Gene fusion detection in 130 pediatric cancer patients. **a** Distribution of 130 pediatric cancer patients colored according to ICD-O-3 primary group. Grey overlay indicates patients for which a clinically relevant fusion was identified by RNA-seq. **b** Schematic overview of the number of fusions at different steps throughout the Fusion-sq pipeline. Subsets discussed extensively in the main text are highlighted in red and available in Table 1. Note that for the 165 high confidence tumor-specific fusions (hcTSFs), 126 occur in patients without a known clinically relevant gene fusion. The same applies to the 27 distinct fusions involving oncogenes or TSGs (onco/TSG). **c** Schematic overview of the Fusion-sq algorithm to find SVs that support fusion predictions by linking the upstream (5’, blue) and downstream (3’, red) partner genes. First, genomic intervals to match RNA-DNA breakpoints are derived based on the intron-exon gene structure and the RNA breakpoints. Next, DNA breakpoints located in these matching intervals are used to identify SVs that link together the partner genes (Methods). **d** Upper panel: schematic representation of high confidence fusion detection based on SVs identified by two or more tools (>50% reciprocal overlap). Lower panel: scatter plot of tumor and normal AF, resulting in classification of fusions in tumor-specific (blue), likely germline (red) and low AF (green). **e** Number of fusions in individual patients: hcTSFs (circles, grouped and color-coded by primary cancer type group) and fusion predictions by RNA-seq with FFPM>0.1 (grey bars). Fusion status is indicated by a black square for clinically relevant fusions and a circle for oncogene/tumor suppressor gene fusions.

### All clinically relevant fusions match to tumor-specific SVs

We first focused on the subset of 30 patients for which a clinically relevant gene fusion was identified by RNA-seq. In all patients, we resolved the tumor-specific SVs for the predicted gene fusions (Table 1). To better understand gene fusions at the genomic level, we investigated the underlying SVs supporting the chimeric transcripts. For all patients but one, the associated duplications (DUP), deletions (DEL) and inversions (INV) were almost identical (>99% overlap) and breakpoints of intra-chromosomal translocations (CTX) either overlapped or were resolved within 13 base pairs (bp) (Fig. 2a). The distances between breakpoints in RNA from the chimeric transcripts and corresponding breakpoints in WGS data from the underlying SVs are highly variable amongst gene fusions and patients (Fig. 2a). Notably, for nine patients the distance between corresponding RNA-DNA breakpoints is larger than 10 kilo base pairs (kb) (Fig. 2a, red lines), which illustrates the advantage of using intron-exon gene structure to define genomic intervals for matching chimeric transcripts with SVs.

**Fig. 2:**
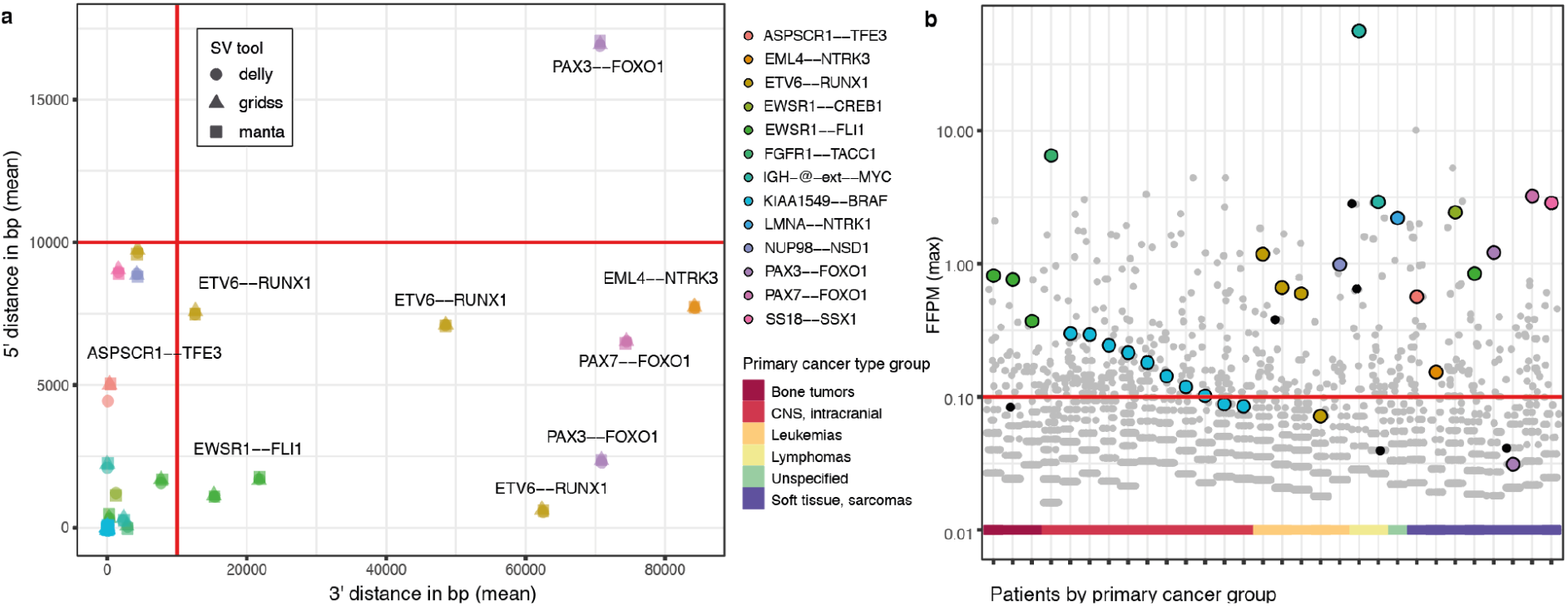
Tumor-specific SVs resolve clinically relevant fusions with high precision and recall. **a** Distance between RNA breakpoints and matching SVs as resolved by DELLY (circle), Manta (square)and GRIDSS (triangle) for the 5’ and 3’ partner genes in 30 patients carrying clinically relevant fusions.. Colors represent specific gene fusions. Red lines indicate 10 kb fixed-size matching intervals and would fail to match SVs for nine gene fusions (labelled). Note that all clinically relevant fusions are detected by at least two SV tools at nucleotide resolution (<10bp, overlapping symbols) except for *ASPSCR1--TFE3*. **b** Gene fusion predictions (all circles) in individual patients (same as in **a**) with their read support in fusion fragments per million total RNA-seq fragments (FFPM). The red line indicates the default cutoff below which fusions are usually discarded (low expression FFPM<0.1). Supporting high confidence tumor-specific SVs were identified for all clinically relevant fusions (colored circles; colors same as in a) and six additional fusions (black) of unknown significance, but not for the remaining RNA-only fusion predictions (grey). Colored bar at the x-axis indicates the primary cancer type group.

One clinically relevant fusion that proved difficult to resolve was *ASPSCR1--TFE3*, because the underlying translocation t(7;X) was identified differently by Manta, DELLY and GRIDSS (Table 1). Manta resolved it as a composite fusion of a CTX + INV in chrX, whilst GRIDSS and DELLY had unusually broad CTX footprints of ∼393 bp and 1181 bp respectively. The associated SV breakpoints of all three tools overlapped SINE mobile elements, which may indicate difficulties in resolving this gene fusion to base pair accuracy in DNA. Despite this potentially complex variant, this translocation could still be mapped to the *ASPSCR1--TFE3* chimeric transcript by Fusion-sq. This enabled further manual inspection of this clinically relevant gene fusion.

The underlying SVs of the other clinically relevant fusions are simple SV events [20] and represent all major SV types, such as deletions, duplications, inversions and translocations. Inter-chromosomal translocations (CTX, 21) were the most common SV type underlying clinically relevant gene fusions as expected[1]. In some cases, these translocations support both the canonical and reciprocal transcripts. As an example, we identified a reciprocal gene fusion product for three out of four patients with an *ETV6--RUNX1* fusion, and for one out of four patients with an *EWSR1--FLI* fusion. In nine patients, we identified duplications resulting in *KIAA1549--BRAF* fusions. Interestingly, the *KIAA1549--BRAF* fusion of patient M218AAA was identified as an inversion by all three tools, and it is likely an inverted duplication as read depth is also increased (+0.41 copy number log2 fold change, CN l2fc). The two remaining clinically relevant fusions are *FGFR1--TACC1* caused by a 420 kb INV, and *LMNA--NTRK1* caused by a 740 kb DEL, showing that a variety of SVs can result in clinically relevant fusion events.

To reduce false positives in RNA-seq fusion detection, filtering based on read support for chimeric transcripts is often implemented with a default minimum of 0.1 fusion fragments per million total RNA-seq fragments (FFPM) [2, 21]. However, by relying only on RNA evidence, four clinically relevant fusions would be missed due to their low expression (Fig. 2b, Table 1), two *KIAA1549--BRAF* fusions (0.08-0.09 FFPM), one *ETV6--RUNX1* (0.07 FFPM) and one *PAX3--FOXO1* (0.03 FFPM). These lowly expressed fusions can be discerned from false positives by integration with WGS. In total, RNA-seq data predicted 3,832 fusions in these 30 patients alone of which 513 passed the read support threshold (FFPM>0.1). In contrast, for 25 patients their clinically relevant gene fusions are the only hcTSFs indicating a high specificity of Fusion-sq. In the remaining five patients, Fusion-sq resolved an additional six hcTSFs with similar support but of unknown significance (Fig. 2b, black dots). This shows that integration of RNA-seq and WGS by Fusion-sq can accurately resolve tumor-specific fusions, effectively removing the need for extensive manual filtering to select which fusions to follow-up by experts.

### Underlying SVs distinguish tumor-specific fusions from healthy chimera

After examining the underlying SVs of known clinically relevant fusions, we returned to the 232 high confidence fusions (hcFs) identified in 86 patients, which are classified into tumor-specific, germline or low allele fraction (Fig. 1b). Both the clinically relevant and additionally detected hcTSFs have similar mean tumor/normal AF of 0.34/0 and 0.28/0 respectively. Supporting their classification, the germline fusions have 0.37/0.36 tumor/normal AF and low AF fusions have 0.02/0 (Additional file 1: Figure S2). While it is counter-intuitive to have high confidence variants with these low AFs, we reasoned that a high number of variant and reference reads could explain this. Indeed, most of these low AF hcFs (20 of 29) originated from amplified regions (3x fold change in read depth, CN l2fc >1.58). Next, we evaluated the efficacy of identifying hcTSFs by 1) assessment of the underlying SV properties, 2) annotation with databases of chimera and SVs, and 3) annotation with cancer-related genes.

Firstly, we investigated whether the underlying SVs of additionally detected gene fusions resemble those of known clinically relevant fusions. Hereto, we mapped the high confidence fusions resolved in individual patients to distinct fusions to account for recurrent fusions. In total the 232 hcFs mapped to 201 distinct fusions of which 145 tumor-specific, 26 germline, 29 low AF, and 1 ambiguous (Table 2, Additional file 1: Figure S3). The fusion labelled as “ambiguous” was categorized differently in two patients and excluded from further analysis. Similar to the clinically relevant fusions, interchromosomal translocations (CTX, 44 of 130) are the most common SV type underlying the additionally identified tumor-specific fusions. The remaining tumor-specific intra-chromosomal SVs are distributed over DUP, DEL and INV (Fig. 3a). In contrast, germline fusions are depleted in CTX events and generally caused by shorter intra-chromosomal SVs. The tumor-specific SVs are ∼27x larger than germline SVs with an average size of 16.2 Mbp compared to 606 kb for germline SVs (Fig. 3b). Only a single germline fusion (*CCDC32--CBX3*) has an underlying CTX, identified with high confidence in five patients and with low-confidence in three patients. *CCDC32--CBX3* is a known healthy chimera and in two of the three low-confidence cases, the SV breakpoints overlap ALU repeat elements which indicates potential mapping difficulties in resolving the breakpoints and suggests a mechanism involved in forming this event. These results show that the types and sizes of SVs underlying tumor-specific gene fusions are distinct from germline events and resemble the SVs of known clinically relevant fusions.

**Fig. 3:**
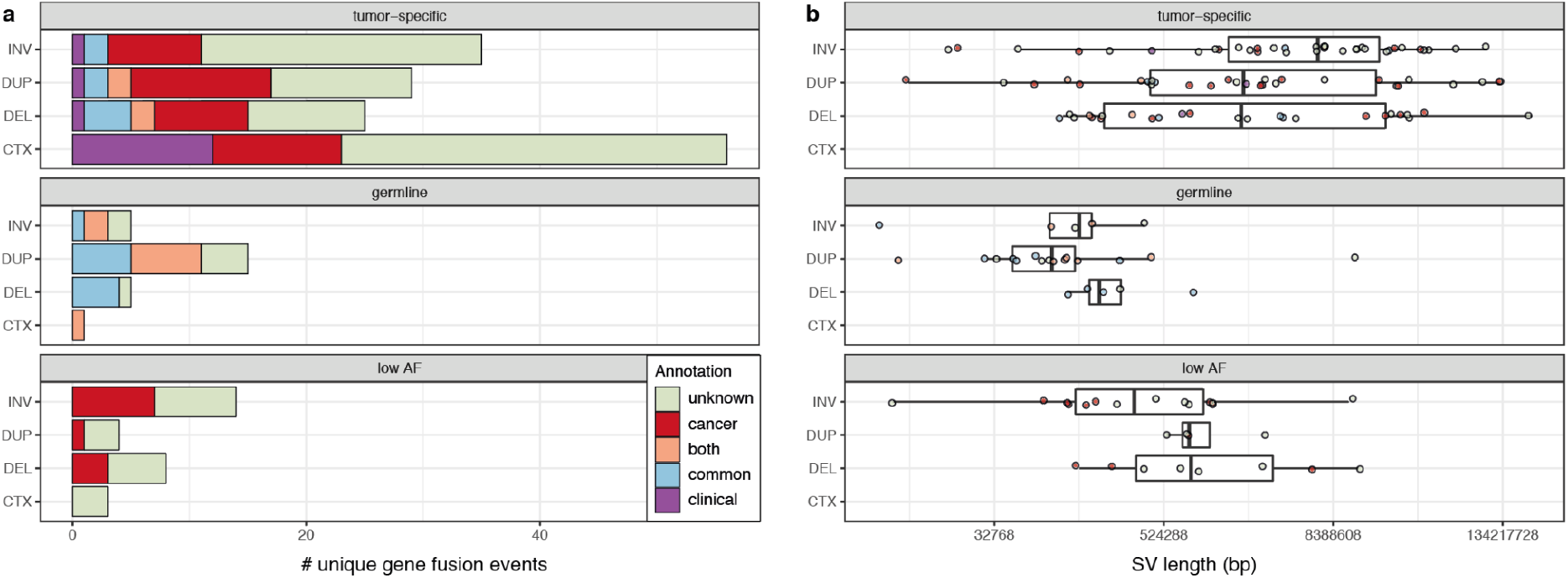
Underlying SVs distinguish tumor-specific fusions from healthy chimera. Number of (**a**) and average SV length in base pairs (bp) for (**b**) distinct high confidence fusions categorized either as tumor-specific (upper panel), likely germline (middle panel) and low AF (lower panel).. Each category is further subdivided according to SV type: inversion (INV), duplication (DUP), deletion (DEL) or intra-chromosomal translocation (CTX). Fusions are colored based on known clinical relevance (clinical; purple), involving a cancer-related gene or cancer chimera (cancer; red), population SV or healthy chimera (common; blue), or both cancer and common (both; orange).

Secondly, to compare hcTSFs with population variants, the gene fusions were annotated against databases cataloguing healthy chimeric transcripts and databases of SVs occurring in the general population as normal variation (Table 2). Overall, nine distinct high confidence fusions occur in a healthy chimera database (4%), which were also all classified as germline fusions. Next, we compared the underlying SVs of fusions to the NCBI Common SV database [22], gnomAD [23] and DGV[24], and found that 28 of the 200 distinct hcFs overlap with SVs occurring in the general population. The majority (16) of these gene fusions were classified as germline and six of those are also flagged as healthy chimera. In total, 73% of the distinct germline hcFs (19 of 26) either overlap a population SV or occur in a healthy chimera database (Fig. 3a). In contrast, only 12 tumor-specific SVs overlap with population SVs (8%) and none occur in a healthy chimera database. These results indicate that the hcTSFs are depleted in germline population variants.

Finally, to identify potentially pathogenic fusions, we compared the results of Fusion-sq with chimera previously detected in cancer and we annotated fusions with cancer-related genes. 44 of the 200 distinct hcFs are present in ChimerDB [25] or the Mitelman database [1]. Most of these are classified as tumor-specific (26), but also germline (9) and low AF (9) fusions occured in these databases. The nine “germline cancer chimera” are additionally annotated as healthy chimera or have underlying intra-chromosomal SVs that overlap with general population SVs (Fig. 3a). This corresponds to previous observations that cancer chimera databases can include passenger fusions [26, 27]. Next, we annotated the fusion partner genes as proto-oncogene or tumor-suppressor gene (TSG). In addition to the known clinically relevant fusions, we identified 29 high confidence fusions involving an oncogene (19) or TSG (19), which are all tumor-specific except for two fusions with a low AF. None of the germline hcFs occur exclusively in a cancer-related resource nor do they affect an oncogene or TSG, further substantiating the classification based on AF of the underlying SVs (Fig. 3a). Annotation of distinct hcFs resulted in identifying 133 of 145 tumor-specific fusions which have no evidence of occurring in the normal population. In summary, we resolved 27 distinct hcTSFs involving an oncogene and/or TSG with similar high confidence as the known clinically relevant fusions, which therefore provide sufficient evidence for further investigation.

### Resolving tumor-specific fusions in individual genomes

Focussing on patients without a known clinically relevant fusion, we identified 126 hcTSFs of unknown significance to investigate further. These fusions are detected in 34 patients across many different cancer types. Generally, each individual patient carries no or only a few hcTSFs (Fig. 1e), whereas a high burden of gene fusions is associated with copy number instability (Additional file 1: Figure S4). Unlike the known clinically relevant fusions that are recurrent within a cancer type, the additionally detected hcTSFs are generally unique to individual patients in this diverse cohort (Additional file 2). Only *MTAP--CDKN2B-AS1* is found in two patients (precursor B-ALL and Pleomorphic xanthoastrocytoma) and this fusion has been previously suggested as potentially pathogenic in melanoma [28]. Therefore, we continued with patient-level analysis to further investigate the functional effects of gene fusions, focussing on fusions involving oncogenes or tumor-suppressor genes.

The benefit of understanding the underlying SVs in a tumor genome is also apparent when resolving fusions in patients showing copy number instability (Fig. 4). High fusion burden (>=7 hcFs, 95th percentile) is associated with a high fraction of genome altered (FGA) by copy number alterations (CNAs) (median 63% vs 3.8%, mean 48% vs 11%, wilcox. p<0.01) and tumor types prone to copy number instability such as osteosarcoma (4x), embryonal sarcoma, embryonal rhabdomyosarcoma, neuroblastoma and ependymoma [29].

**Fig. 4:**
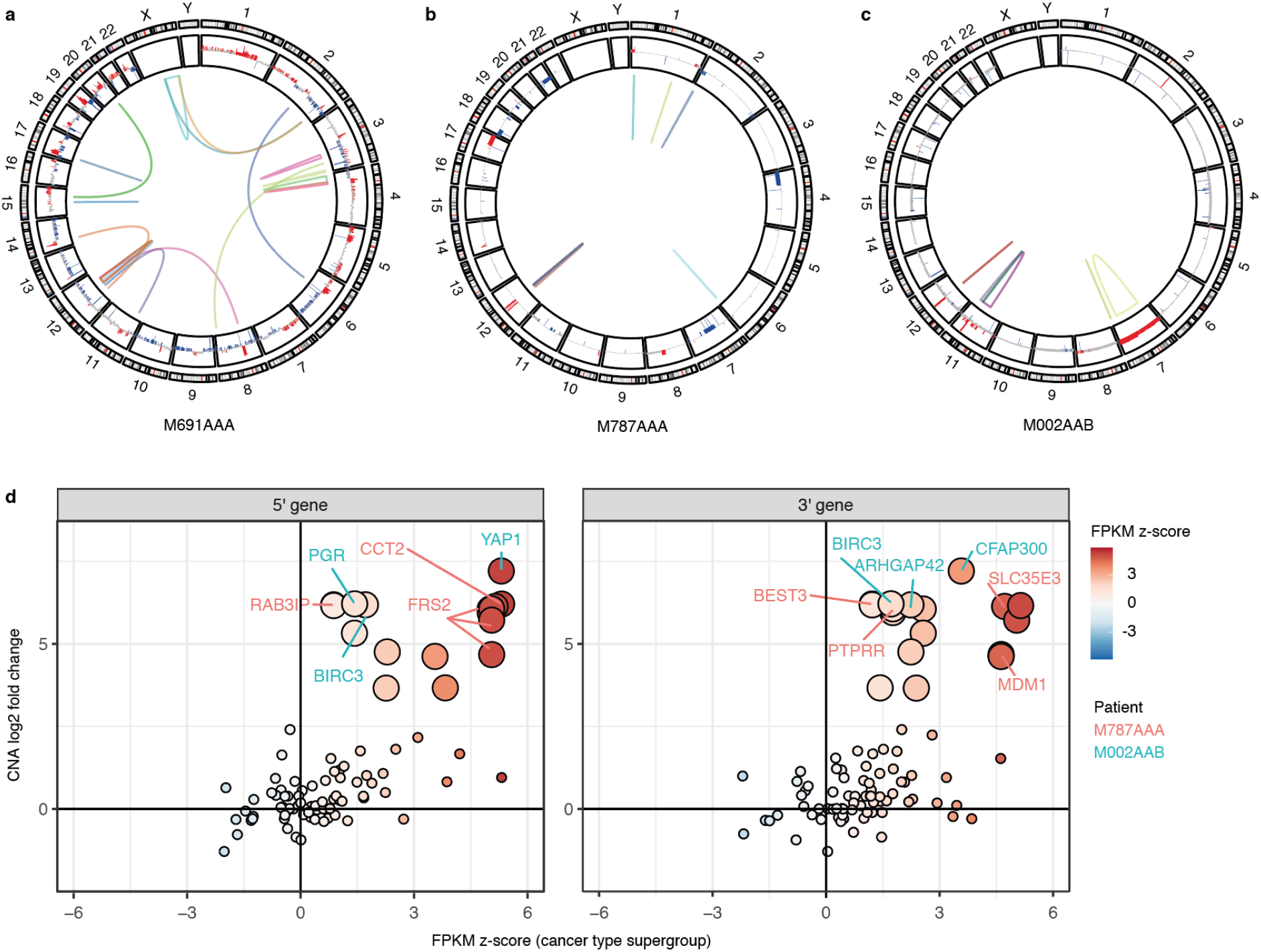
Resolving tumor-specific fusions in patients with high copy number instability and focal amplifications. Circos plots of osteosarcoma patient M691AAA (**a**), neuroblastoma patient M787AAA (**b**) and embryonal rhabdomyosarcoma patient M002AAB (**c**). The plots are annotated with high confidence tumor-specific gene fusions (multi-colored links) and copy number gains (red) and losses (blue). **d** Relationship between gene expression changes of the 5’ (left panel) and 3’ (right panel) partner gene relative to the cancer type supergroup (FPKM z-score, color scale displayed on the right) and copy number (CN) log-2 fold changes in read depth. Genes located inside amplified regions (CN l2fc>1.58) are displayed with larger circles. Cancer-related genes are labelled according to their patient of origin.

To further investigate this association between copy number instability and high fusion burden, we closely studied four osteosarcoma patients carrying *TP53* and *ATRX* fusions (Additional file 1: Figure S5). These patients’ tumors have many dispersed CNAs (median FGA 67%) and also many hcFs distributed across their genomes (range 7-19, Fig. 1e, Additional file 1: Figure S5a). In three of these osteosarcoma patients, we resolved fusions involving the first exon of *TP53* and in each case a different downstream (3’) partner gene resulting from an inversion and two translocations; t(17;6), t(17;20) (Additional file 1: Figure S5b) (Table 1). Translocations with the first exon of *TP53* have previously been identified as cancer driver events in osteosarcoma [30]. Therefore, these fusions are potentially also pathogenic in these patients, especially since a driver mutation had yet to be identified for these tumors. Apart from the *TP53* gene fusions and associated SVs, we did not observe additional CN losses or deleterious SNVs in the *TP53* loci of these patients. In the fourth osteosarcoma patient (M691AAA, Fig. 4a), we detected an *ATRX* fusion which has been suggested previously as a potential driver mutation for osteosarcoma [30]. For the *ATRX* fusion, gene expression concomitantly was reduced relative to the cancer type supergroup (−3.3 z-score FPKM (zfpkm), p<0.01). Similarly, the group of patients with a *TP53* fusion showed reduced expression relative to the cancer type supergroup (0.88 vs 2.0 log2 FPKM, p<0.05) (Additional file 1: Figure S5c). However, this was not clear for the individual patients, illustrating that the underlying SV provides additional evidence for a disruptive SV event that could not be easily derived from RNA-seq alone.

Multiple fusions originating from highly amplified regions were resolved in two patients with neuroblastoma (M787AAA, Fig. 4b) and embryonal rhabdomyosarcoma (M002AAB, Fig. 4c). The neuroblastoma patient has a focal amplification in chromosome 12q13-15 involving the oncogenes *MDM2* and *CDK4* (4-6 copy number log2 fold change, CN l2fc). In this region, we identified 13 high confidence, low AF gene fusions. Including fusions previously identified as cancer-related chimera (i.e. *FRS2--MDM1, FRS2--PTPRR, CCT2--BEST3, RAB3IP--BEST3*) of which both partner genes are overexpressed due to the amplification (Fig. 4d). Similarly, patient M002AAB carries seven fusions originating from a focal amplification in chr11q22 (5-7 CN l2fc) which are detected with high confidence but some with a low AF. Here also, the fusion partner genes are overexpressed which is consistent with the amplification (Fig. 4d). Although chr11q22 amplification itself is not known to be clinically relevant, we resolved multiple fusion combinations with oncogenes *BIRC3, PGR* and in particular *YAP1. YAP1--CFAP300* involves exons 1-5 of the *YAP1* oncogene which is highly amplified (7.2 CN l2fc) and overexpressed (5.3 zfpkm, p<0.1). Also, this exon 1-5 fragment has previously been identified as pathogenic when fused to other 3’ partner genes by activating the TEAD pathway [31, 32].

These case studies illustrate that Fusion-sq can confidently resolve fusions in unstable genomes with a high FGA or complex alterations. For some tumor-specific fusions, the underlying SVs are potentially pathogenic (e.g. the *TP53* and *ATRX* fusions) while in other cases the fusions seem to be the result of copy number instability. In addition, gene fusions in amplified regions can exhibit high expression of their partner genes due to the underlying CN gain. Therefore, characteristics of underlying SVs are key for interpreting potential functional effects of individual gene fusions, especially in patients with unstable genomes.

### Tumor-specific fusions affecting gene expression

Gene fusions can activate oncogenes or disrupt tumor-suppressor genes (TSG) and the resulting pathogenic effects can be reflected in dysregulation of gene expression. To identify potentially pathogenic fusions, we aimed to assess the functional effects of the 27 distinct hcTSFs involving an oncogene or TSG (in **bold**) (Table 1). In addition, we included two composite fusions that were detected by both GRIDSS and DELLY but differed in how they reported the underlying SV (Table 1). For *LINC01344--****TERT***, the translocation breakpoints overlapped but reported in addition either an insertion or a duplication in *TERT*. Also for ***ETV6****--IGL*, the same translocation was resolved, but the additional inversion in *IGL* had different breakpoints. The SVs underlying the other 27 hcTFs involving oncogenes or TSGs represent all the simple SV types: CTX (10), DUP(5), DEL(7) and INV(5). As a proxy for functional effect, we combined expression data, underlying SVs and gene annotation.

Overexpression of oncogenes resulting from gene fusions is often suggested as an activation mechanism [2]. However we did not observe an enrichment of oncogenes amongst overexpressed fusion partner genes (>1.96 zfpkm) compared to the cancer type supergroup or the full cohort. Instead, our data suggests an association with copy number gain (CN l2fc>0.58) irrespective of gene annotation, as reflected by about half of the fusions with CN gain found to be overexpressed (20 of 42). Also, partner genes of hcTSFs are more often affected by CN gain and overexpression together, than each occurring separately (2.3x odds ratio, p<0.05). Although a cohort wide pattern is lacking, we identified fusions in individual patients for which the 3’ oncogenes are significantly overexpressed relative to the cancer type supergroup (p<0.1): ***PAX3****--****WWRT1*** (2.1 and 3.0 zfpkm), *LINC01344--****TERT*** (2.4 zfpkm) and *SYMPK--****MEF2B*** (2.1 zfpkm) (Fig. 5). In addition, we identified a *MED14--****HOXA9*** fusion and associated overexpression of *HOXA9* (2.2 zfpkm, p=0.15) in a pre-T-cell lymphoblastic leukemia patient (M385AAA). In this case, *HOXA9* overexpression did not reach significance because an acute myeloid leukemia patient (M975AAA) with a *NUP98--NSD1* fusion, which is known to dysregulate *HOXA* genes, also has a high *HOXA9* expression (3.0 zfpkm, Additional file 1: Figure S6) [9]. Taken together, these results indicate that resolving the SV and copy number can help to distinguish between overexpression due to underlying amplifications and expression changes resulting from the fusion of a 3’ partner gene to an active upstream promoter.

**Fig. 5:**
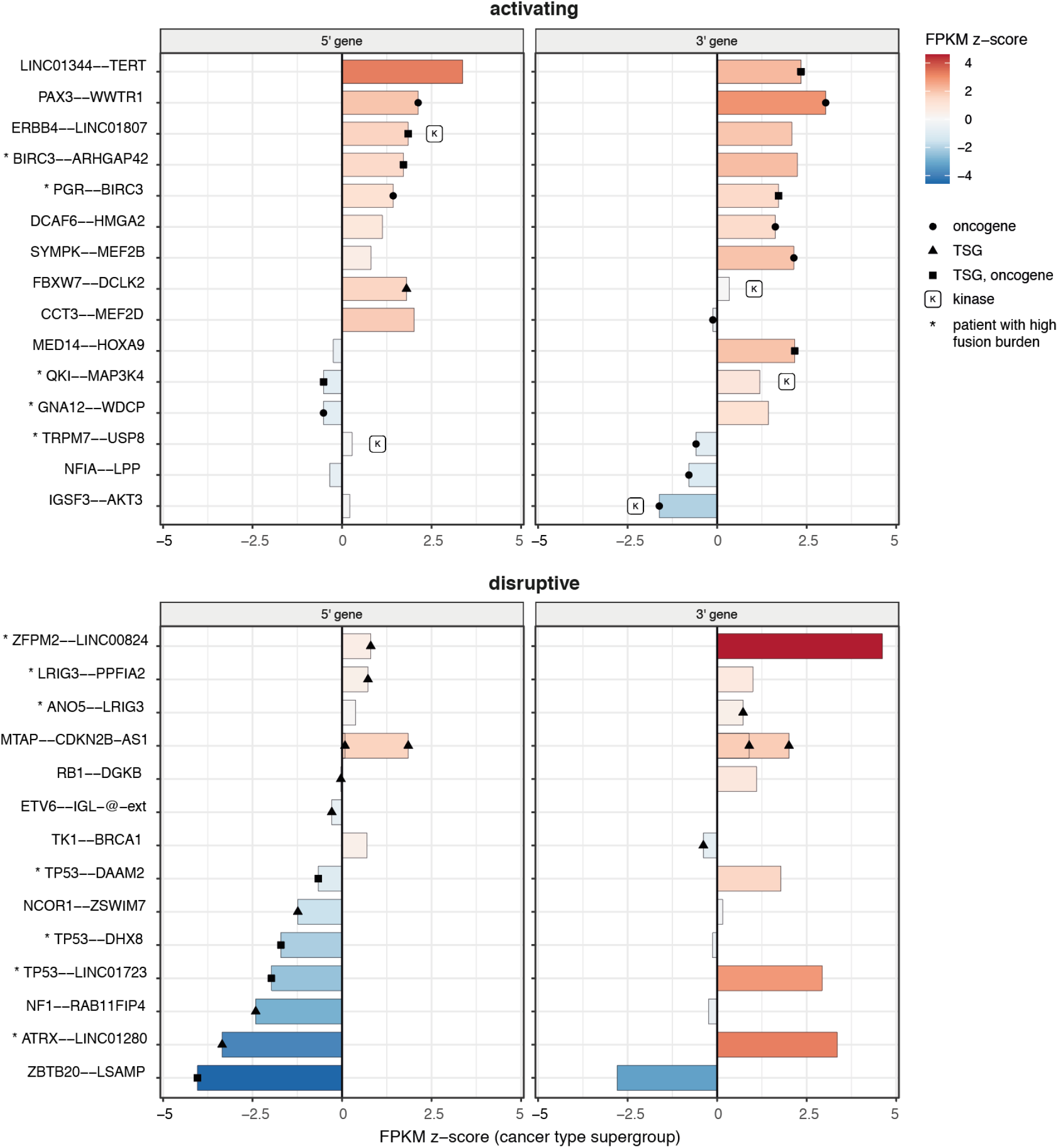
Fusion partner gene expression can indicate oncogene activation or tumor-suppressor gene disruption. Gene expression changes for the 5’ (left) and 3’ (right) partner genes of tumor-specific gene fusions involving oncogenes and/or tumor-suppressor genes (TSG). Changes are relative to the cancer type supergroup (FPKM z-score; color scale on the right). Individual 5’ or 3’ genes are marked as either oncogene (circle), TSG (triangle), both TSG and oncogene (square) or kinase (k). Fusions were divided into ‘activating’ or ‘disruptive’ functional effect categories based on partner gene annotation, and marked with an asterisk if they originate from patients with a high gene fusion burden.

Gene fusions can also be pathogenic through activation of kinases due to e.g. loss of an auto-inhibitory domain or increased dimerisation. We identified six fusions involving kinases, of which three in patients with unstable genomes (*QKI--MAPK3K4, TRPM7--USP8, STK40--OSCP1*). In patients without known driver alterations, we identified *ERBB4--LINC01807, FBXW7--DCLK2* and *IGSF3--AKT3*. However, the relevance of these fusions remains unclear. The *ERBB4* fusion arises from a tandem duplication and is associated with an increase in gene expression (1.9 zfpkm, pval=0.1) in a region with reduced read depth (−0.13 CN l2fc), but the *ERBB4* kinase domain is not involved in the gene fusion. Similarly, the underlying SV of the *AKT3* fusion also showed only the last exon is included in the fusion product and there are no known mechanistic links for *DCLK2* activation in nephroblastoma. Combining the evidence from SVs, CN and gene expression suggests that these fusions are unlikely to result in kinase activation.

### Fusions disrupting tumor-suppressor genes

Gene fusions that involve tumor-suppressor genes (TSGs) are potentially pathogenic, as the underlying SVs can disrupt the TSGs. Similar to oncogenes, we did not find a general trend of downregulation in the 19 fusions involving TSGs, however SVs intersecting a gene can be disruptive regardless of whether gene expression is affected. Three specific cases did show a significant decrease in gene expression (p<0.1) relative to the cancer type supergroup: ***ZBTB20****--LSAMP* (−4.0 zfpkm, p<0.05), ***NF1****--RAB11FIP4* (−2.4 zfpkm) and the previously mentioned ***ATRX****--LINC01280* (−3.5 zfpkm) in a patient with osteosarcoma. Both *ZBTB20--LSAMP* and *NF1--RAB11FIP4* have been previously detected in other adult cancer types [1, 25]. In addition, *ZBTB20* and *NF1* play important roles as TSG in pediatric cancers and *LSAMP* is suggested as a potential TSG in osteosarcoma and neuroblastoma [33–35].

To investigate whether TSGs could be disrupted by gene fusions without this being reflected in gene expression changes, we sought to identify co-occurring somatic single nucleotide variants (SNVs). We found co-occurring fusions and SNVs for *RB1* and *NCOR1* that could indicate double-hits of these TSGs, despite the fact that no reduction in gene expression was observed for *RB1* (−0.1 zfpkm) and only a minor reduction for *NCOR1* (−1.2 zfpkm). In an adrenal cortical carcinoma patient (M152AAD), the *RB1* gene was affected by a translocation resulting in the *RB1--DGKB* fusion and by a splice donor site mutation (chr13:48473390_GGTGA/G, tumor AF 0.32, normal AF 0, Additional file 1: Figure S7a). In a second patient with pre B-cell lymphoblastic leukemia (M930AAB), we identified a 158 kb deletion in the *NCOR1* locus resulting in the fusion *NCOR1--ZWIM7*, as well as a frameshift mutation (chr17:16070282_C/CT, tumor AF 0.85, normal AF 0.03, Additional file 1: Figure S7b). This deletion overlaps a healthy population SV and is likely ALU-mediated given that the breakpoints fall in ALU repeats. However, in this case, the deletion is specific to the patient’s tumor sample (0.33 tumor AF, 0 normal AF). Also, for both *RB1* and *NCOR1*, gene fusions with different 3’ partner genes have been previously identified in cancer [1, 25]. In combination, the co-occurrence of deleterious SNVs and the promiscuity of these fusions seem to suggest that the presence of these gene fusions is indicative of TSG disruption.

## Discussion

Discovery of novel driver gene fusions in pediatric cancer has been severely limited by a lack of methods for genome-wide unbiased detection. To systematically discover tumor-specific gene fusions, we developed Fusion-sq which integrates chimeric transcripts from RNA-seq with SVs from WGS using intron-exon gene structure. Previous studies combining RNA-seq and WGS data for gene fusion detection have reported a large variation in validation rates, which is likely due to differences in the detection and integration methods used [14, 27, 36, 37]. We identified supporting SVs for ∼2.5% of (unfiltered) gene fusion predictions, which is similar to the previously reported 91% false positive rate for STAR-Fusion predictions [36]. In contrast, the PCAWG transcriptomics group identified supporting SVs for 82% of RNA alterations in their study [14]. This difference is likely due to stringent pre-filtering of gene fusion predictions and lenient RNA-DNA breakpoint matching with 500kb intervals. In contrast, we identified SVs that precisely support chimeric transcripts and abstained from expression-based filtering, which would otherwise have excluded four clinically relevant fusions with low expression levels. The recently published tool MAVIS also integrates SVs and RNA-seq, but treats them separately and requires recurrence to identify gene fusions [37]. Other pediatric cancer studies focusing on precision oncology rely heavily on experts to perform the data-integration manually [16, 38] thereby limiting its applications for candidate discovery.

A common strategy to identify potentially pathogenic fusions and distinguish them from passenger fusions is to focus on recurrent events [14, 16, 37]. This approach yielded little results in our pediatric pan-cancer cohort, likely due to the relatively small size, high heterogeneity of cancer types and low mutation burden of pediatric cancers. In addition to the known clinically relevant fusions that we successfully identified, the only other recurrent fusion we found is *MTAP--CDKN2B-AS1*. Yet, we were able to identify multiple potentially pathogenic gene fusions in individual patients by leveraging SV properties and gene expression data. We showed that analyzing the underlying SVs of gene fusions can help to discern tumor-specific fusions from likely passenger fusions that are the result of normal transcription processes, germline SVs or are related to copy number instability. Consistent with reports from adult cancers [1, 27], we found associations between high gene fusion burden and a high FGA, or unstable regions such as focal amplifications. Although it was suggested that this relationship might be absent in pediatric cancers[29], the observed discrepancy with our analysis could be due to how copy number instability is quantified or differences in cohort composition rather than a mechanistic difference between adult and pediatric cancers. Therefore, our results suggest that alternative strategies to recurrence can identify potentially pathogenic fusions. Recurrence is nevertheless an important criteria for defining fusions of clinical relevance and is an important rationale for further expanding genomic studies.

Detection of gene fusion chimeric transcripts using RNA-seq data is limited to actively transcribed genes, and fusions with non-coding elements may be missed. In addition to protein-coding fusion products, SVs that displace an enhancer can cause unusually high expression of oncogenes[3]. However, these “enhancer hijacking” events fall outside the scope of this study as they lack chimeric transcript evidence. For example, an *IGH--MYC* translocation was identified with FISH in patient M879AAA and in WGS data we also identified the underlying reciprocal translocation from the *IGH* locus to ∼200kb upstream of *MYC*. However, this translocation results in an enhancer exchange and not in the formation of a chimeric transcript, hence it could not be identified by Fusion-sq. Also chimera resulting from intergenic fusions can be missed, since they arise from SVs followed by additional splicing alterations and may not have SV breakpoints corresponding to their chimeric transcript [39]. Moreover, requiring orthogonal support from WGS is highly effective in filtering potential false positives from RNA-seq, as well as other chimeric transcripts without underlying genomic mutations that result from normal transcription processes (i.e. read-through events or cis/trans splicing) [12, 26]. Many of these RNA-only chimeric transcripts also occur in healthy tissues and are less likely to have a pathogenic effect than fusions caused by tumor-specific SVs[12][13]. Our results show distinct differences between types and sizes of SVs underlying tumor-specific and germline high confidence fusions (hcFs) (Fig. 3). The majority of tumor-specific hcFs were found to be the result of inter-chromosomal translocations. These properties provide a strong basis for distinguishing passenger gene fusions from potentially pathogenic variants based on WGS SV data [27].

In addition to the high confidence subset of 232 gene fusions, we also identified 33 fusions for which two or more tools resolved underlying SVs but with differences in SV type and exact breakpoint location. This is likely the result of mapping difficulties due to repeat elements at the breakpoints. We found evidence of composite fusions having two underlying SVs, for example the known clinically relevant fusion *ASPSCR1--TFE3* (Table 1) which was reported as t(17;X) in combination with an inversion. In addition, we identified two composite fusions *LINC01344--TERT* and *ETV6--IGL* both of which have an underlying translocation combined with a duplication likely mediated by segmental duplications surrounding the breakpoints. As a result, they are not in the high confidence set as their underlying SVs are identified with slight differences between SV tools. Despite these challenges, we were able to identify gene fusions with high confidence also for patients with highly rearranged genomes and focal amplifications. Further analysis of both DNA and RNA with long-read sequencing would be beneficial to fully resolve underlying complex SVs and gene fusions in detail [40].

Overall, we identified 27 distinct gene fusions in 19 patients that involve oncogenes or TSGs and display similar characteristics to fusions that have previously been linked to tumor etiology. For some patients, these candidate fusions add to the list of variants of unknown significance which have been identified in their tumor genomes, but for others the identified gene fusion presents a strong candidate for follow-up studies.

In case patients have copy number unstable genomes and carry many gene fusions, the pathogenicity of individual fusions is difficult to assess. For example, the co-amplification and resulting overexpression of *MDM2/CDK4/FRS2* in neuroblastoma patient M787AAA is clinically relevant [41], and the fusions originating from this amplification are more likely to be passenger events [27, 42]. This shows that resolving the underlying SVs and copy number alterations (CNAs) can help to distinguish expression changes due to “catastrophic” genomic events from pathogenic fusions where 3’ (onco)genes are upregulated due to fusion with an active promoter. However, we did not observe strong cohort-level trends of gene fusions resulting in 3’ oncogene overexpression, likely because of the highly specific associations between oncogenes and certain pediatric cancer types. Instead, we found an association between CN gain and fusion partner gene overexpression, consistent with observations in adult cancers that CNAs are the main contributing factor to gene expression changes[14].

For a subset of patients, we resolved fusions that potentially activate 3’ oncogenes as reflected in expression changes. For example, we identified a fusion associated with *TERT* overexpression in a patient diagnosed with liver cell adenoma (M637AAB, Additional file 1: Figure S8a). This fusion warrants further investigation, because *TERT* activation may be a factor contributing to malignant transformation to liver carcinoma [43]. Also, in a yolk sac tumor (M014AAA), we identified a gene fusion with the *ERBB4* kinase, which was previously found to be overexpressed in yolk sac germ cell tumors [44] and suggested as a potential drug target [45]. Although the kinase domain is not involved in this patient’s gene fusion (Additional file: Figure S8b), also kinase-dead mutants can have functional consequences in cancer [46].

In addition, we identified two gene fusions involving transcription factors that potentially result in gain-of-function chimeric proteins. We identified a *PAX3--WWRT1* fusion in an embryonal rhabdomyosarcoma patient (M911AAA) resulting from a translocation t(2;3) (tumor AF 0.81) which gives rise to a fusion product involving exons 1-7 of *PAX3*. Canonical *PAX3--FOXO1* fusions involve the same *PAX3* exons, which suggests that *PAX3--WWRT1* may have similar functional effects as the *PAX3/7* driver gene fusions in alveolar rhabdomyosarcoma (Additional file: Figure S8c). In a patient with pre-T-cell lymphoblastic leukemia (M385AAA), we identified *MED14--HOXA9* as in-frame gene fusion involving the homeobox transcription factor domain (tumor AF 0.47, normal AF 0.06, Additional file 1: Figure S8d). While this specific fusion has not yet been reported, diverse mechanisms of *HOXA9* activation might impact treatment and prognosis in T-cell leukemia [47, 48]. Notably, a *FLT3-ITD* was identified in this patient too, and concurrent *FLT3-ITD* and *HOXA9* overexpression was suggested to be potentially pathogenic in acute myeloid leukemia [49]. Taken together, these examples highlight candidate gene fusions which may activate oncogenes relevant in these cancer types and warrant further investigation of their pathogenic potential and relevance for the clinic.

Disruption of TSGs through gene fusion events has been previously reported in pediatric cancer, but the impact of this mechanism remains underestimated [50]. While gene fusions can result in activation of oncogenes, less is known about fusions involving TSGs [2]. These fusions that potentially disrupt TSGs can be promiscuous in their partner genes and breakpoints which makes them difficult to detect with targeted assays [7, 8]. For example, the three *TP53* fusions occurred with different partner genes and gene expression data alone did not clearly indicate a disruptive effect. However, osteosarcoma samples can have low tumor cell percentages that complicate gene expression analysis [30]. In these patients, resolving the underlying SVs provided additional evidence supporting potential *TP53* disruption.

In total, we identified three cases where TSG fusion partners are downregulated in expression: *ATRX--LINC01280, ZBTB20--LSAMP and NF1--RAB11FIP4*. The *ATRX* fusion is potentially pathogenic in osteosarcoma since it has been suggested as a driver before [30]. Both *ZBTB20--LSAMP* and *NF1--RAB11FIP4* involve known TSGs in neurological pediatric cancers and have been reported in adult cancers [1, 25]. However, they have not yet been described as markers of TSG disruption in pediatric cancers. In a neuroblastoma patient (M909AAA), we resolved a *ZBTB20--LSAMP* fusion due to a 1 Mb duplication which indicates a potentially pathogenic disruptive event (Additional file 1: Figure S8e). *ZBTB20* is associated with neuronal differentiation and disruption of this process is a known oncogenic factor in neuroblastoma [51]. The neural cell-adhesion protein *LSAMP* has been suggested as a potential TSG in neuroblastoma [33, 35] as well as other cancer types such as osteosarcoma [34], however the mechanism is less clear. Both *ZBTB20* and *LSAMP* are downregulated and for *ZBTB20* low expression is correlated with poor prognosis in a publicly available neuroblastoma dataset (Additional file 1: Figure S9) [51].

The *NF1--RAB11FIP4* fusion identified in a patient with Pleomorphic xanthoastrocytoma (M535AAA), was caused by a 185 kb deletion with a disruptive effect on *NF1* (Additional file 1: Figure S8f). *NF1--RAB11FIP4* fusions have been previously detected in multiple cancer types [25]. *NF1* is also an important TSG that is recurrently mutated in pediatric CNS tumors [52] and in rare instances also specifically xanthoastrocytoma [53]. Of note, germline *NF1* alterations in combination with a second-hit somatic mutation can indicate sensitivity to immunotherapy [52]. Although we only identified a somatic deletion with 0.5 tumor AF, the expression analysis provides additional evidence that the *NF1* gene is significantly disrupted in this patient.

Finally, TSG disruption can also occur through alternative mechanisms as with the *MTAP--CDKN2B-AS1* fusions. These were the only recurrent fusions in our cohort and have been previously reported in melanoma [28]. The presence of a *MTAP--CDKN2B-AS1* fusion indicates disruption of the *CDKN2A* locus via two possibly parallel mechanisms; directly resulting from the deletions in the *CDKN2A* locus causing the gene fusion, and indirectly through upregulation of *CDKN2B-AS1* (0.9-2.0 zfpkm, Additional file 1: Figure S10) which can have a repressive effect via Polycomb or RNA interference [28, 54]. Therefore, presence of *MTAP--CDKN2B-AS1* fusions can indicate concurrent *CDKN2A* disruption on multiple regulation levels. Taken together, these examples illustrate that resolving underlying SVs can provide crucial orthogonal support and additional evidence for TSG disruption, thereby facilitating mechanistic understanding and clinical interpretation.

## Conclusion

To increase our understanding of pediatric cancer, new approaches should be developed to identify novel driver gene fusions. In this paper, we identified tumor-specific gene fusions with high confidence by combining chimeric transcripts from RNA-seq and SVs from WGS data. Resolving the underlying SVs enables confident detection of known clinically relevant fusions, as well as discovery of potentially pathogenic fusions by distinguishing them from artefacts and healthy-occurring events.

SVs can aid selection of tumor-specific fusions for targeted therapies and aid minimal residual disease monitoring by providing allele fractions and exact breakpoints. Furthermore, we identified 27 potentially pathogenic tumor-specific gene fusions involving oncogenes and tumor-suppressor genes and demonstrated how these events can be linked to gene expression changes. Regardless of the pathogenicity of the fusion itself, it can be a marker for underlying genomic rearrangements especially in the case of TSG disruption. Rare gene fusions present an interpretation challenge and, without recurrence, they require further investigation into biological mechanisms or pathways. For example, through the integration of expression data and in combination with the mutational landscape of the patient. The approach used in this study is not only useful for pediatric cancer, but can also be applied in adult cancer for identifying candidate pathogenic fusions. Overall, we show the power of integrating RNA-seq gene fusion predictions with WGS structural variants, which can aid discovery and interpretation of pathogenic fusions for precision oncology applications.

## Materials and methods

### Sample preparation and sequencing

Data was collected as part of the biobanking initiative at the Princes Máxima Center for Pediatric Oncology, and resulted in a pan-cancer cohort of 130 patients. The inclusion criteria used were: the availability of informed consent, paired tumor-normal sequencing WGS data and RNA-seq data of the tumor of sufficient quality (see quality control metrics), and the sample being representative of the cancer type group (i.e. presence of tumor material in the sample).

Following the institute’s standardized biobanking protocols (Hehir-kwa, 2021, manuscript under revision), RNA and DNA were isolated from fresh frozen tumor tissue and as a matching normal, DNA was isolated from whole blood. Blood and bone marrow samples were enriched for monocytic cells using Ficoll. Total RNA was isolated from tumor samples using the AllPrep DNA/RNA/Protein Mini Kit (QIAGEN) according to standard protocol on the QiaCube (Qiagen) RNA-sequencing (RNA-seq) libraries were generated from 300ng RNA using the KAPA RNA HyperPrep Kit with RiboErase (Roche) and sequenced with NovaSeq 6000 (2×150 bp) (Illumina). DNA was isolated from paired tumor-normal samples also using the AllPrep DNA/RNA/Protein Mini kit. Whole-genome sequencing (WGS) libraries were generated from 150 ng DNA using the KAPA DNA HyperPlus kit and NovaSeq 6000 sequencing platform (Illumina).

### RNA and WGS sequencing data pre-processing

Pre-processing of RNA-seq and WGS was done with the institute’s standardized pipelines implementing GATK 4.0 best practices for variant calling using a wdl and cromwell-based workflow [55, 56].

Data quality was assessed with Fastqc (version 0.11.5) to calculate the number of sequencing reads[12]. Picard (version 2.20.1) for both WGS and RNA metrics output such as insert size and MarkDuplicates [13]. The RNA sequencing reads were aligned using Star (version 2.7.2b) to GRCh38 and gencode version 31 [14]. WGS reads were aligned using BWA mem (0.7.13) to GRCh38.

Quality control (QC) metrics are available for all samples (Additional file 3). For WGS, a minimum median coverage of 25x for normal samples and 60x for tumor samples was used. The percentage of duplicate reads was reasonable as well with median 8% and maximum 13% for tumor samples, 7% and 13% for normal samples respectively. Patient M129AAA was resequenced once to achieve sufficient coverage.

For RNA data, a minimum unique reads count of 30M was used based on the Picard total reads and percentage of duplicates.

### Diagnostic process

As part of the institute’s routine diagnostic process, patients were diagnosed according to ICD-O-3 guidelines combining histopathological and molecular characteristics by a pathologist and molecular tumor board. In order to achieve sufficient sample size for some of the downstream analyses, the ICD-O-3 primary cancer type groups were further grouped into three cancer type supergroups: hemato, neuro and solid. Hemato contains the leukemia (1) and lymphoma (2) primary groups, Neuro the CNS tumors (3) and neuroblastomas (4). Note that there were no patients with retinoblastoma (5) in our cohort. The solid group is composed of all other primary cancer groups (6-12).

### Variant calling

Gene fusion predictions were obtained from tumor RNA-seq using STAR Fusion (version 1.8.0) [57] and GRChr38/Gencode v31 CTAT Oct 2019. Fusion predictions involving human leukocyte antigens or mitochondrial genes are filtered out, but no other pre-filtering based on RNA support was done.

Single nucleotide variants (SNVs) were inferred from paired tumor-normal WGS by Mutect2 from GATK 4.1 [58] and pathogenicity was predicted by variant effect predictor (VEP) (version 92)[59], according to the GATK4 standards. Somatic SNVs were filtered based on tumor variant allele frequency (AF) > 0.05 and predicted impact (MODERATE or HIGH). Somatic copy number alterations (CNAs) were identified with the GATK4 pipeline according to their standards. We generated a panel of normals (PON) from 18 normal samples prepared and sequenced under the same conditions and used this for normalisation. The allelic imbalance ratios were calculated using 1000 genomes, autosomal SNP sites with a minor allele frequency (MAF) > 0.1.

Structural variants (SVs) were inferred from paired tumor-normal WGS using Manta [17] (version 1.6), DELLY [18] (version 0.8.1) and GRIDSS [19] (version 2.7.2). Due to technical issues with running the tool, no GRIDSS output was available for four patients (M863AAC, M479AAA, M156AAA, M606AAA). SVs were not filtered based on quality, read support or somatic/germline annotation. Since these tools vary in how they classify SVs as somatic or germline, we performed this classification based on variant allele fraction (AF) of the paired tumor and normal samples.

The calculation of variant AF was done in agreement with the developer’s recommendations for every tool. In the case of Manta, the tool outputs separate files with “somatic SVs” for the tumor and “diploid SVs” for the normal sample. The tumor and normal AF was calculated for the variants in the somatic file using the number of spanning read pairs and split reads that strongly (Q30) support the reference or variant alleles. Variant AF = SRV+PRV / (SRR + PRR + SRV + PRV). For the variants scored under the diploid assumption, only normal AF could be calculated.

For DELLY and GRIDSS the AF was calculated for all variants. DELLY recommends reference/variant allele supporting pairs for imprecise variants (AF = DV/(DR+DV)) and reads for precise variants (AF = RV/(RR+RV)). GRIDSS recommends using the supporting fragments (VF) that combine split reads, discordant pairs and assembly-based support. (AF = VF / (VF + REF + REFPAIR) for variants larger than the max fragment size distribution, and excluding the REFPAIR for smaller variants (<1000bp).

### Fusion-sq algorithm

#### Prepare matching intervals

Gene fusion predictions from RNA-seq were used to derive genomic intervals for SV breakpoint matching taking into account intron-exon gene structure. First, transcripts were retrieved from the ENSEMBL database based on ENSEMBL gene stable identifiers provided by STAR-Fusion, or based on genomic location in case an identifier was lacking (e.g. immunoglobulin genes). Second, a hierarchy of matching intervals was generated, from more to less precise: 1) intron adjacent to the RNA breakpoint, 2) alternative splice junction +/-10 bp from breakpoint, 3) flanking interval spanning +/-500bp at each side of the breakpoint which is subsetted to the gene body if available, 4) RNA breakpoint to start/end of the gene body. The adjacent intron interval can differ between transcripts, therefore the union of introns was used for initial matching of RNA and SV breakpoints. The transcript-specific intervals later used to match SV breakpoints to individual transcripts.

#### Match SVs to gene fusions

SVs identified by Manta, DELLY and GRIDSS were matched to gene fusion predictions based on these genomic intervals. Note that respecting the hierarchy of the intervals is important because of the inherent overlap between the intervals, i.e. introns fall inside the gene body. We conclude that fusions are validated by WGS if SVs are found that link the up- and downstream (5’/3’) partner genes by any of these genomic intervals.

If no SV was identified that directly links the 5’ and 3’ gene, an attempt was made to resolve the fusion by a composite of two SVs. All SV breakpoints originating in respectively the 5’ and 3’ partner gene’s adjacent intron/flanking/splice-junction intervals were considered. Fusions were flagged as ‘composite’ when two SVs were identified that respectively originate from the 5’/3’ partner genes and have their “other end” in close proximity (5kb) therefore indirectly but effectively linking the partner genes to form a fusion.

#### Combine supporting SVs

As the final step of the pipeline, the supporting SVs from the different tools were integrated for the WGS validated fusions. Each fusion was annotated with the genomic breakpoints and SV characteristics (i.e. SV type, size, tumor and normal allele fractions, breakpoint quality filters). Also, the SV breakpoints were used to select corresponding transcripts for the partner genes and annotated with the involved exons and gene fragments based on this selection. This step also further annotates the precision of SV support by distinguishing between SVs that link the 5’/3’ partner genes via introns of individual transcripts *(gup/gdw_location = intron)* and SVs that link via the union of introns used during matching but do not satisfy this strict criterium for both partner genes (*gup/gdw_location = intron_consensus)*. In some cases, multiple fusion predictions were validated by the same SV and subsequently considered as a single fusion. Vice versa, multiple SVs could be linked to a single fusion prediction as well.

We further selected high confidence fusions (hcFs) based on the support by at least two SV tools and the location of the SV breakpoints relative to the chimeric transcript. Fusions were labeled as ‘*precise_location*’ in case the SVs link partner genes by their adjacent introns, flanking regions or alternative splice junctions. To be considered as *‘high_confidence’*, these supporting SVs had to be resolved as the same event by at least two tools based on 50% reciprocal overlap and matching SV type. SV breakpoints smaller than 30bp were resized to 30bp during matching. In case multiple high confidence SVs support the fusion, the SV with the highest tumor AF is selected.

Fusions were classified based on AF after resolving the underlying high confidence SVs, since this filters out potential additional lower confidence SVs. Fusions were classified as tumor-specific, (likely) germline and low AF based on mean tumor and normal AF of the associated SVs.

- Tumor specific: (tumor AF - normal AF) > 0.05 & (tumor-normal)/normal ratio > 1.5
- Germline: normal AF > 0.05 & (tumor-normal)/normal ratio < 1.1
- Low AF: (tumor AF - normal AF)<0.05 & normal AF <0.05

### Expression data generation and analysis

Gene expression was analyzed using featureCounts from Rsubread (version 1.32.4) with Gencode v31 CTAT Oct 2019 annotation and settings *allowMultiOverlap=T, largestOverlap=T* and *countMultiMappingReads=F*.

Gene expression alterations were assessed with z-scores of log2 transformed gene length normalized read counts (Fragments Per Kilobase of transcript per Million mapped reads, FPKM). Expression values were first log transformed, after which the group mean and standard deviation were calculated. z-*score = (fpkm - fpkm_mean)/ fpkm_sd*. Gene expression z-scores (zfpkm) were reported relative to the full cohort, cancer type supergroup and the primary cancer group. Normality of the log-transformed FPKMs was assessed with Shapiro for each group separately to assess the validity of using gene expression z-scores for outlier analysis. As threshold for aberrantly expressed genes, +/-1.96 z-score was used corresponding to 95% confidence interval (p<0.05). To account for cases where gene expression is not normally distributed in a certain group, we also assessed whether a patient carrying a gene fusion has a significantly different gene expression than other patients in their subgroup using Wilcoxon rank sum test.

To study whether fusions with oncogenes or tumor-suppressor genes as partner genes are associated with gene expression changes, we assessed subset enrichments with Fisher’s exact test and compared zfpkm distribution amongst subsets of fusions with Wilcoxon rank sum tests.

Overexpression is defined as >1.96 zfpkm relative to the cancer type supergroup. Distributions of zfpkm supergroup scores were compared relative to all tumor-specific high confidence gene fusions (hcTSFs) for fusions with and without downstream oncogenes. Similar analysis was conducted for downregulation of tumor-suppressor genes. Next, we investigated whether expression changes and copy number (CN) changes were associated. Hereto again the Fisher’s exact test was used for subset enrichments relative to all hcTSFs and Wilcoxon rank sum tests for changes in zfpkm distributions. CN gain was defined as >1.58 read depth log2 fold change which corresponds to 3x fold change.

### SV type and size analysis

SV properties were analyzed for the high confidence gene fusion that have underlying SVs supported by at least two SV tools. To account for differences between how these tools report events, SV properties were harmonized between tools prior to analysis. SV breakpoints were classified into the major types of simple SVs based on their relative orientation: deletions, duplications, inversions and inter-chromosomal translocations. Sizes of SVs were regarded as positive numbers and only considered for intra-chromosomal events. One gene fusion *(AC063944*.*1--LINC00882*) was resolved as tumor-specific in one patient and as germline in another, therefore it was labeled as ambiguous and ignored during this analysis.

### Recurrence analysis

The number of unique occurrences of fusions across patients was used during recurrence analysis. Every occurrence of an upstream-downstream (5’/3’) partner gene pair in a patient is counted once, ignoring multiple predicted breakpoints in a single patient. Fusion directionality was respected, so canonical and reciprocal gene fusions were regarded as two distinct events.

### Integration of CN data and SVs

Copy number (CN) segments were mapped to SVs based on genomic location. To analyze read depth of SVs, we considered the CN ratio log2 fold change (l2fc) of SV breakpoints and calculated a weighted average from overlapping segments.

### Measures of copy number instability

Fraction of genome altered (FGA) was calculated relative to the expected autosomal genome size, excluding chromosome X and alternate loci: 2875001522 bp.

*FGA = number of base pairs >0*.*2 absolute CN l2fc / expected autosomal genome size*.

In addition to the FGA, the maximum CN l2fc was used to gauge whether patients had focal amplifications.

The subset “patients with a high gene fusion burden” is defined by the 95th percentile (7 or more) of high confidence gene fusions classified as either tumor-specific or low AF. Patients in this subset: M809AAA, M479AAA, M691AAA, M002AAB, M787AAA, M040AAA, M606AAC, M597AAC.

### Annotation of SVs

To verify how SVs link together the fusion partner genes independently of RNA-seq evidence, SVs were annotated with introns based on overlap with canonical transcripts. For each partner gene, a canonical transcript was selected based on stepwise filtering until a single transcript remained: MANE select, the tags “basic, CCDS, APRIS”, protein coding, transcript support level and coding sequence length. The transcript annotation was retrieved from Gencode v31 [14].

As an additional confirmation of the classification between tumor-specific and germline SVs based on AF, SVs underlying gene fusions were compared to SVs occurring in the general population. Hereto, SVs were retrieved from NCBI Curated Common Structural Variants (nstd186) [22], gnomAD Structural Variants (nstd166) [23] from NCBI repository and from DGV [24] (version 2020-02-25) accessed on 2021-03-11. SVs supporting gene fusions were matched to population SVs based on 50% reciprocal overlap, regardless of SV types as variant type annotation differs per database and SV detection method. Fusions were flagged in case their underlying SVs matched to population SVs from any of these databases (*anno_sv_population*) (corresponding column of Table 1).

Furthermore, SVs were annotated with repeats and segmental duplications to assess whether their breakpoints reside in rearrangement-prone genomic regions. Hereto, repeats and segmental duplications tracks were retrieved from UCSC table browser accessed on 2021-04-20 [60]. Repeats from RepeatMasker were pre-filtered by repeat class (LINE, SINE, LTR) and completeness (<50 bp of repeats left) to prevent spurious annotations. Gene fusions were annotated with identifiers of repeats and segmental duplications overlapping the underlying SV start/end coordinates (*repeat_family, segdup)*.

### Annotation of gene fusions

To identify whether gene fusions were previously reported in either healthy tissue or cancer samples, we compared our findings to chimeric transcript databases. Fusions were annotated as healthy chimera based on the default annotation from STAR-Fusion [57]. For the annotation of cancer chimera, we used ChimerDB 4.0 (retrieved on 2021-02-17) [25] and the Mitelman database (v20201015, retrieved on 2021-01-07) [1] matching exact gene pairs. *(anno_cancer_chimera)*.

To aid the interpretation of gene fusions and select potentially pathogenic gene fusions, we assigned gene-level properties to the 5’/3’ partner genes of fusions based on their stable ENSEMBL identifiers and/or gene names. Cancer-related gene datasets were retrieved from COSMIC [61] (cancer gene census v92), OncoKB (accessed on 2021-04-14) [62] and Grobner [29]. For COSMIC and OncoKB we adopted their annotation of oncogenes and tumor-suppressor genes (TSGs). As a pediatric cancer resource, we retrieved recurrently mutated genes identified by Grobner *et al*. and used “amplification” as proxy for oncogene and “deletion/gene-disrupting structural variant” for TSG. Similarly, genes and fusions are annotated as kinase based on the human kinome [63] (retrieved from www.kinase.com on 2021-01-16). For each annotation, it was specified whether the fusion partner gene has that property (*gup_label, gdw_label)*, and the annotation was summarized on the level of the gene fusion for easier selection (*anno_has_onco_or_tsg, anno_has_kinase)*. Finally, gene fusions were also annotated with cytobands retrieved from the UCSC table browser on 2021-05-06.

To summarize annotations for visualisation and reporting (including in Figure 3 and Tables 1 and 2), we annotated fusions *(annotation)* based on known clinical relevance (clinical), involving a cancer-related gene or cancer chimera (cancer), population SV or healthy chimera (common), or both cancer and common (both). For further analysis, we selected potentially pathogenic fusions based on whether they contain an oncogene or TSG.

### Correlating gene expression and prognosis

Kaplan Meier plots were retrieved from the R2 genomics analysis and visualisation platform. (http://r2platform.com/, accessed on 2021-07-12). Results were obtained with KaplanScan, which calculates the optimum gene expression threshold for survival analysis with statistical testing. Data from neuroblastoma samples in the publicly available Versteeg dataset was used for this analysis http://www.ncbi.nlm.nih.gov/geo/query/acc.cgi?acc=GSE16476

### Gene fusion schematics

Gene fusion schematics were generated with Protein Paint from the St. Jude cloud (https://proteinpaint.stjude.org/[64, 65] using the default RefSeq transcripts in hg38 and genomic coordinates of the underlying SVs resolved with Fusion-sq.

## Supporting information

Table 1 column key

Additional file 2: Recurrent gene fusions

Additional file 3: Quality Control metrics

## Declarations

### Ethics approval and consent to participate

Informed consent has been obtained for all subjects involved in this study through the Máxima biobank informed consent procedure and corresponding protocol. The Máxima biobank protocol has been approved by the Medical Ethics Committee of the Erasmus Medical Center in Rotterdam, The Netherlands, under reference number MEC-2016-739. Approval for use of the subject’s data within the context of this study has been granted by the Máxima biobank and data access committee (https://research.prinsesmaximacentrum.nl/en/core-facilities/biobank-clinical-research-committee), biobank request nr. PMCLAB2018.017.

## Consent for publication

Not applicable

## Availability of data and materials

The WGS and RNA-seq datasets supporting the conclusions of this article are available in the EGA repository. Code for fusion-sq is available through https://github.com/princessmaximacenter/fusion-sq.

Extended data tables are included as supplemental data.

## Authors’ contributions

P.K., J.H.K., F.C.P.H. and B.B.J.T. substantially contributed to the conception and design of the study. I.A.E.M.B., P.K. and J.H.K. drafted the article. I.A.E.M.B performed data curation, normalization, computational analyses and generated the figures. Sequencing experiments were performed by M.vT and E.S. C.C, S.B, A.J, E.W, D.vdL, H.K were involved in analysis of the NGS data and management of metadata. J.J.M, J.M, J.D, L.K and W.C.P provided input on variant interpretation. All authors discussed the concepts and contributed to the final manuscript.

## Acknowledgements

We would like to thank all children and their families for participating in our research. We would also like to acknowledge all clinicians, lab technicians and biobank committee members for acquiring patient consent, collecting and processing of samples as well as general biobanking logistics. We also thank lab members for fruitful discussions and feedback.

## Figures and tables

Table 1: Gene fusions for which underlying SVs are resolved with high confidence as tumor-specific or low AF variants. Also included are the three gene fusions caused by a composite SV detected by at least two tools: *ASPSCR1--TFE3, ETV6--IGL-@-ext, LINC01344--TERT*. For convenient subset viewing, use the *anno_clinically_relevant* and *anno_has_onco_or_tsg* flags.

Table 2: Gene fusions from Table 1 mapped to distinct fusions such that fusions occurring in multiple patients are merged and listed only once.

