## Additional file 1 for "Systematic discovery of gene fusions in pediatric cancer by integrating RNA-seq and WGS"

### Supplementary data

**Table of contents:**


**Additional file 1**

- Figure S1: Gene fusions supported by one or more SV tools
- Figure S2: Allele fraction of distinct fusions per SV type
- Figure S3: Filtering predicted gene fusions to a high confidence subset
- Figure S4: High gene fusion burden is associated with copy number instability
- Figure S5: Osteosarcomas with a *TP53* gene fusion
- Figure S6: *HOXA9* gene expression and gene fusion status
- Figure S7: Co-occurring fusions and SNVs indicative of TSG disruption
- Figure S8: Potentially pathogenic gene fusion candidates in individual patients
- Figure S9: Association between *ZBTB20* gene expression and survival.
- Figure S10: *MTAP--CDKN2B-AS1* fusions and associated expression changes of *CDKNB-AS1* and *CDKN2A*

**Additional file 2: Recurrent gene fusions**

**Additional file 3: Quality control metrics**

### Additional file 1

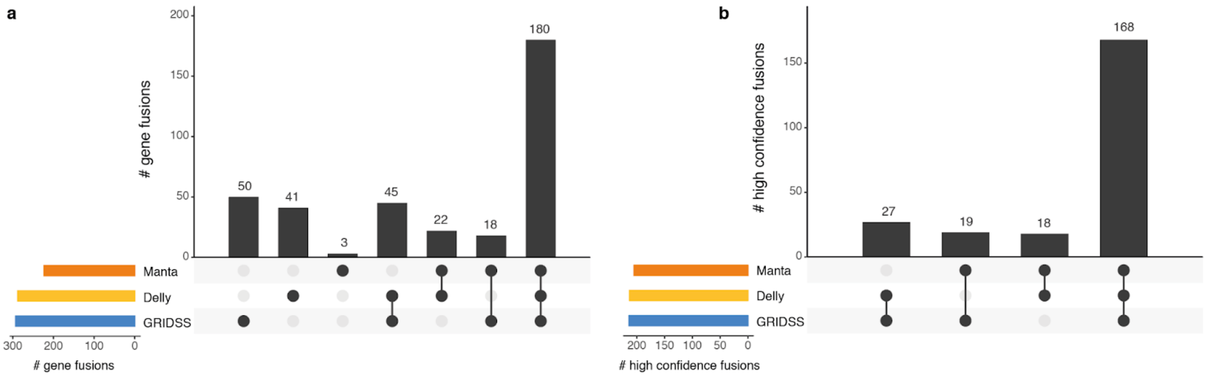


**Figure S1: Gene fusions supported by one or more SV tools**
**a** Distribution of gene fusions supported by SVs detected by Manta, DELLY and/or GRIDSS.
**b** same as **a** but for the high confidence subset.
NB: no GRIDSS output was available for four patients (M863AAC, M479AAA, M156AAA, M606AAA)

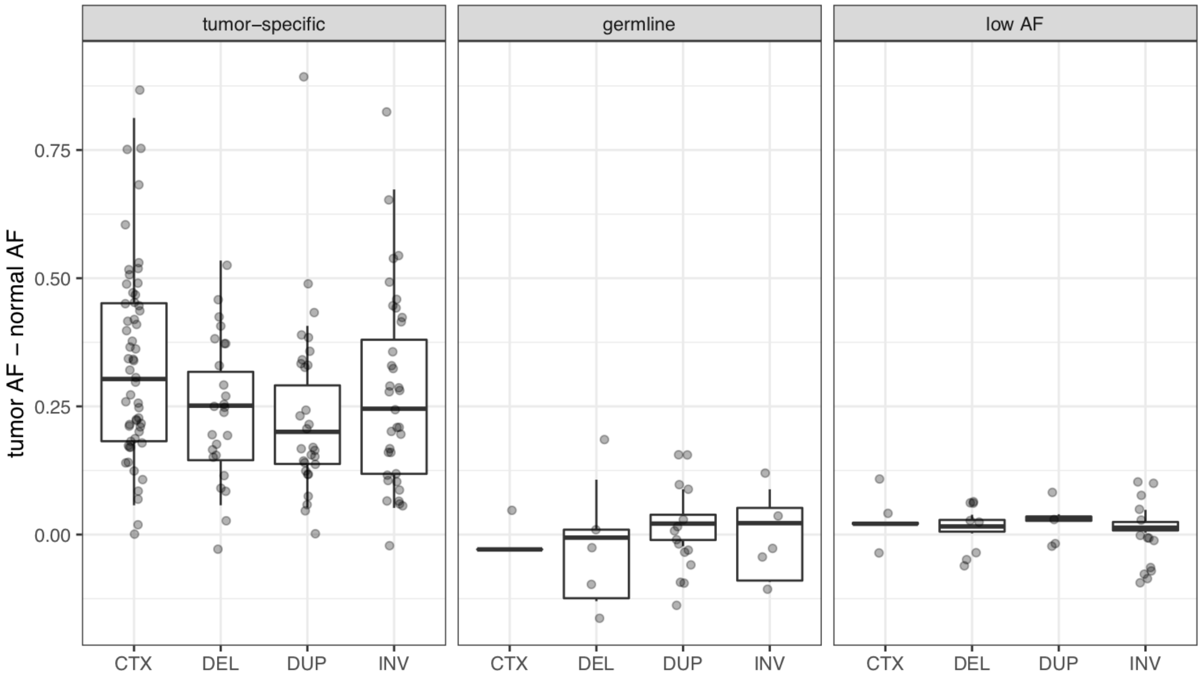


**Figure S2: Allele fraction of distinct fusions per SV type.**
Allele fractions (AF) of distinct fusions are shown for either tumor-specific (left), likely germline (middle) or low AF (right).

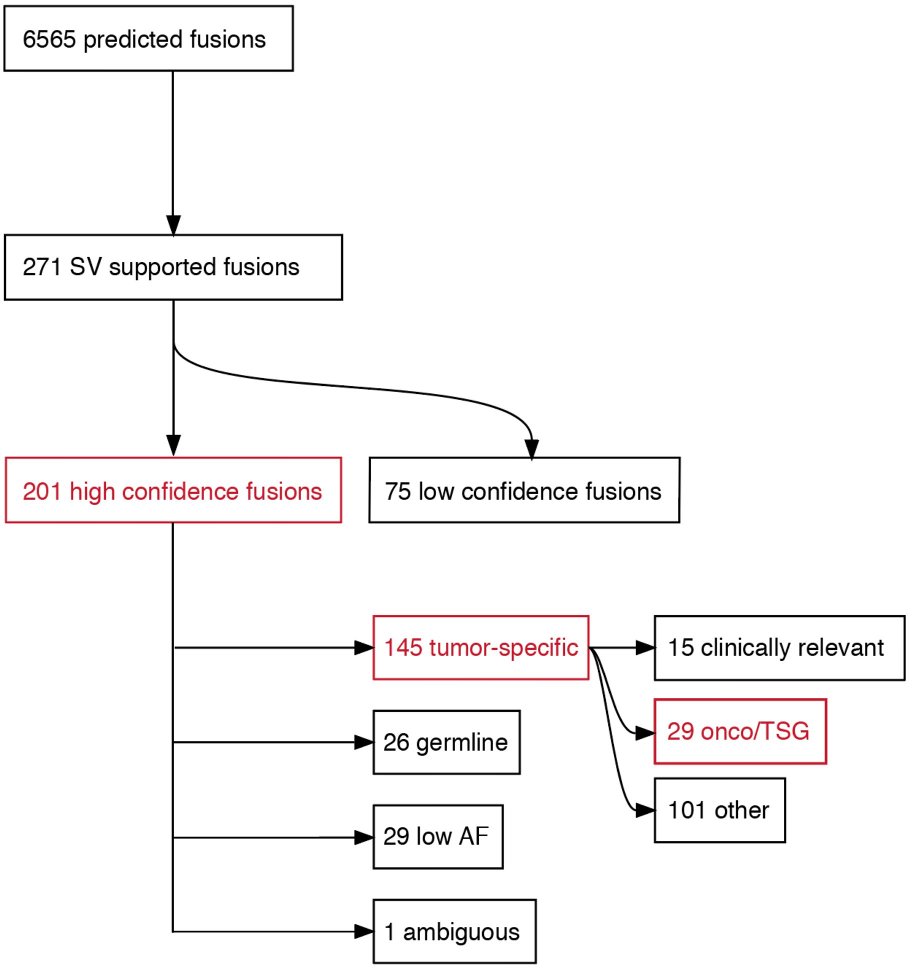


**Figure S3: Filtering predicted gene fusions to a high confidence subset**
Schematic overview of the number of distinct fusions throughout the Fusion-sq pipeline. The steps are the same as in Fig. 1b, but gene fusions are mapped to “distinct fusions” such that fusions occurring in multiple patients are merged and counted only once.
Fusions were labelled “ambiguous” and excluded in case they were categorized differently amongst patients. Subsets discussed in the main text are highlighted in red and available in Table 2.

Note that the 27 distinct fusions involving an oncogene or tumor-suppressor gene (onco/TSG) in patients in the main text specifically refers to those identified in patients without known clinically relevant gene fusion. The other 2 out of 29 fusions are detected in patients carrying a known clinically relevant gene fusion and therefore not discussed in the main text.

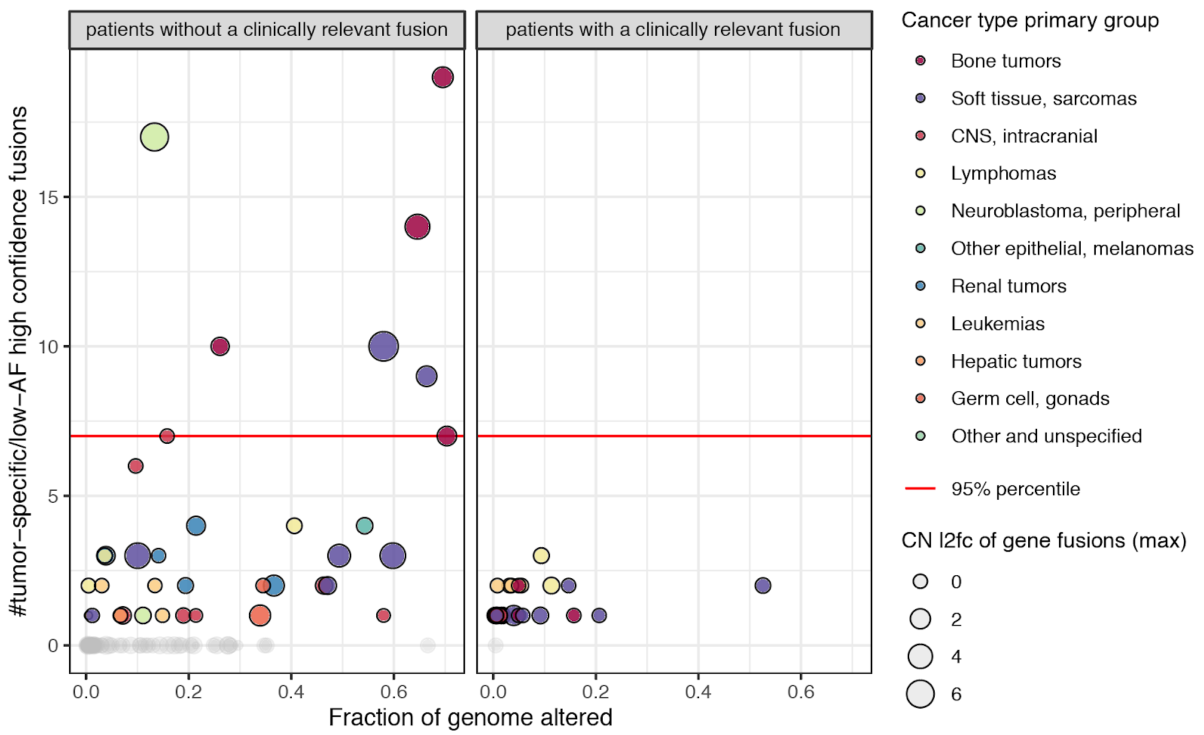


**Figure S4: High gene fusion burden is associated with copy number instability.**
Relationship between the fraction of genome altered by copy number alterations (FGA) and the number of high confidence gene fusions classified as tumor-specific or low AF. Colors represent primary cancer type groups, circle size corresponds to the maximum copy number log2 fold change (CN l2fc) underlying the gene fusion.
Red line indicates 95% percentile. Patients equal or higher are labeled as high gene fusion burden (Methods) and have either a high FGA or gene fusions arising from focal amplifications (high CN l2fc).

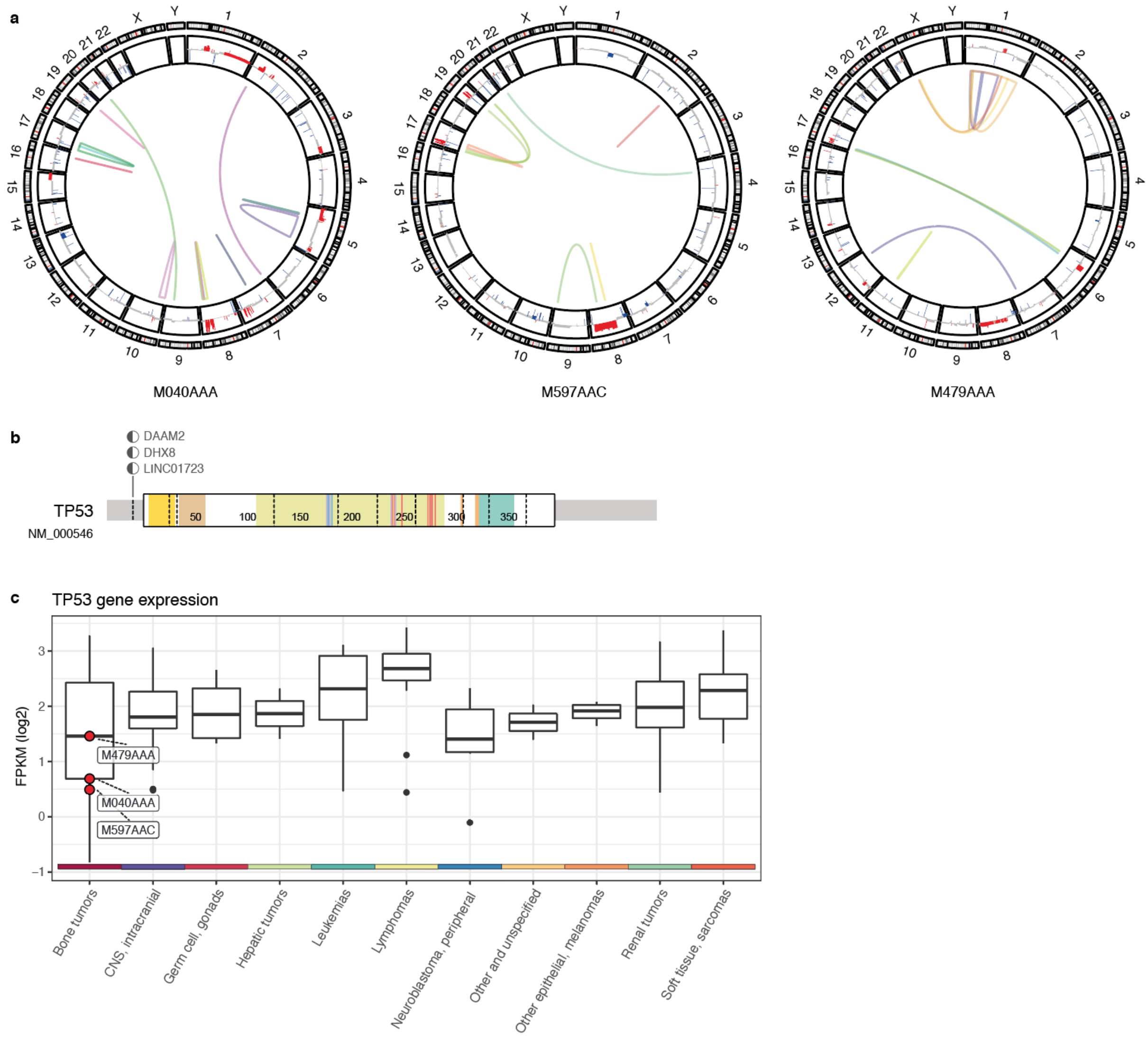


**Figure S5: Osteosarcomas with *TP53* gene fusion.**
**a** Circos plots for patients carrying a *TP53* gene fusion (M479AAA, M597AAC, M040AAA) with high confidence tumor-specific gene fusions (multi-colored links), copy number gains (red) and losses (blue). **b** Schematic representation of the resolved gene fusions with *TP53* exon 1 and downstream (3') partner genes. **c** Gene expression levels of *TP53* (log2 FPKM) of patients carrying a *TP53* gene fusion (red circles) in comparison to all patients split according to their primary cancer type group (violin plots). Colored bar at the x-axis indicates the primary cancer type group.

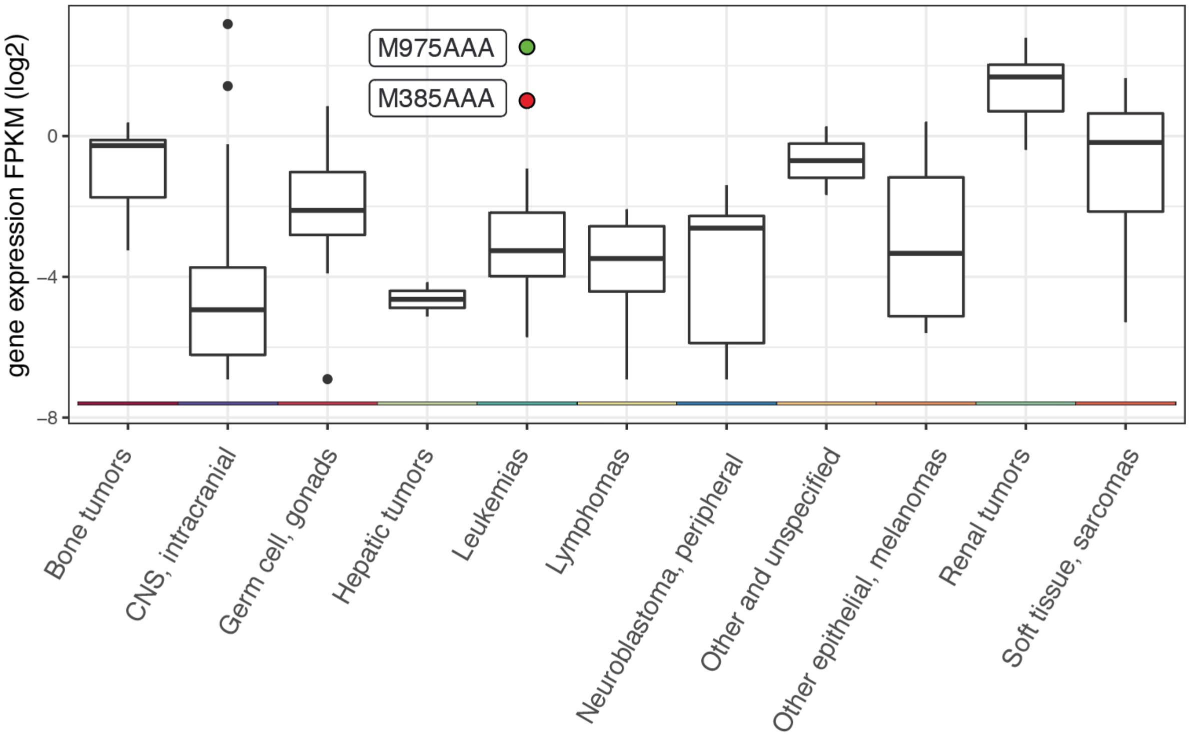


**Figure S6: *HOXA9* gene expression and gene fusion status**
Gene expression levels of *HOXA9* (log2 FPKM) for patient M385AAA carrying a *MED14--HOXA9* gene fusion (red circle), patient M975AAA with a *NUP98--NSD1* gene fusion (green circle) and all patients split according to their primary cancer type group (violin plots).

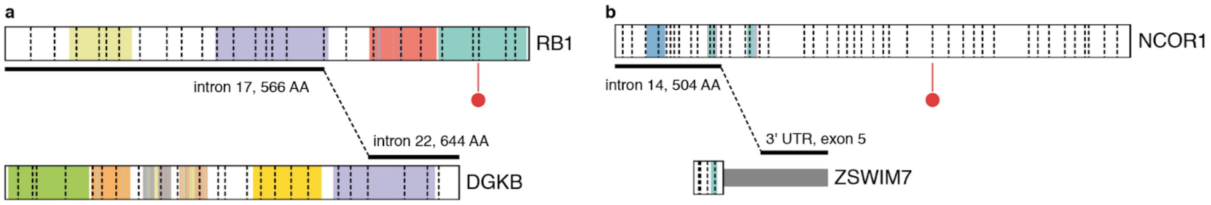


**Figure S7: Co-occurring fusions and SNVs indicative of TSG disruption**
Schematic representation of gene fusions and somatic SNVs (red flags) detected in the same patient for **a** *RB1--DGKB* in patient M152AAD and a splice donor site mutation (chr13:48473390_GGTGA/G) and for **b** *NCOR1--ZSWIM7* in patient M930AAB and a frameshift mutation (chr17:16070282_C/CT)

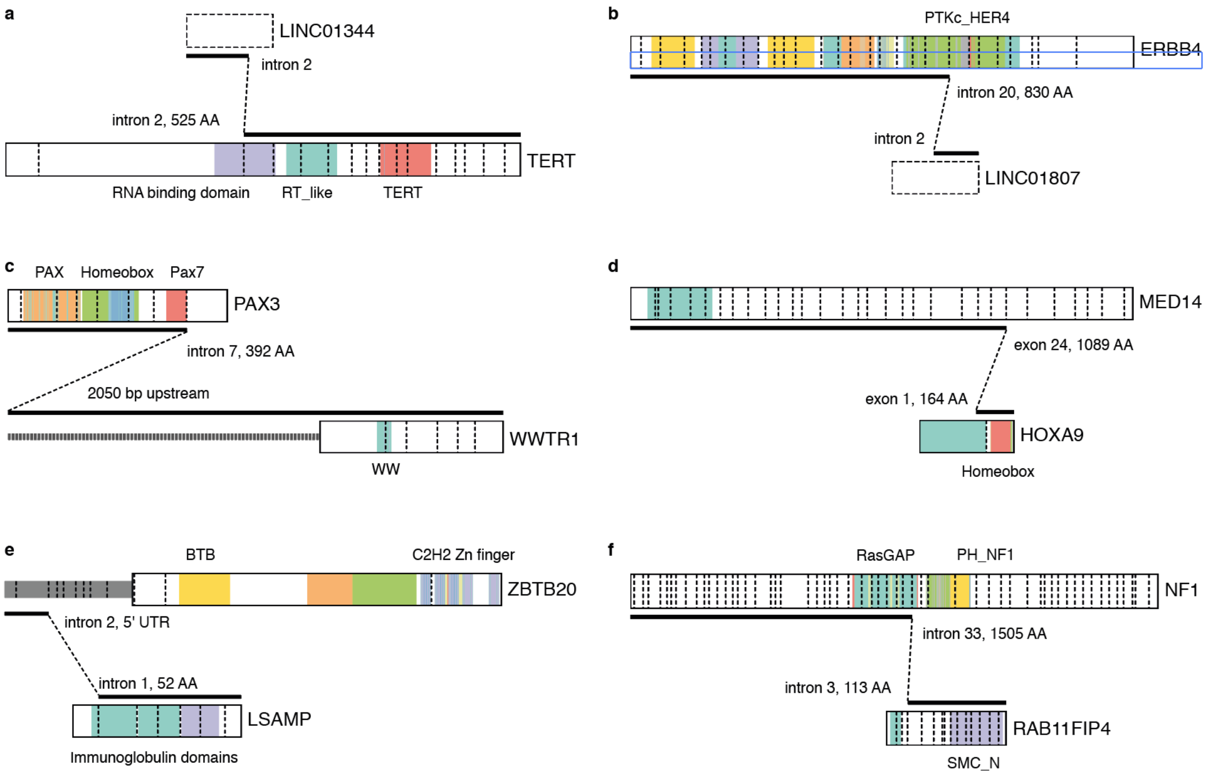


**Figure S8: Potentially pathogenic gene fusion candidates in individual patients**
Schematic representations of gene fusion *LINC01344--TERT* (**a**), *ERBB4--LINC01807* (**b**), *PAX3--WWRT1* (**c**), *MED14--HOXA9* (**d**), *ZBTB20--LSAMP* (**e**) and *NF1--RAB11FIP4* (**f**).

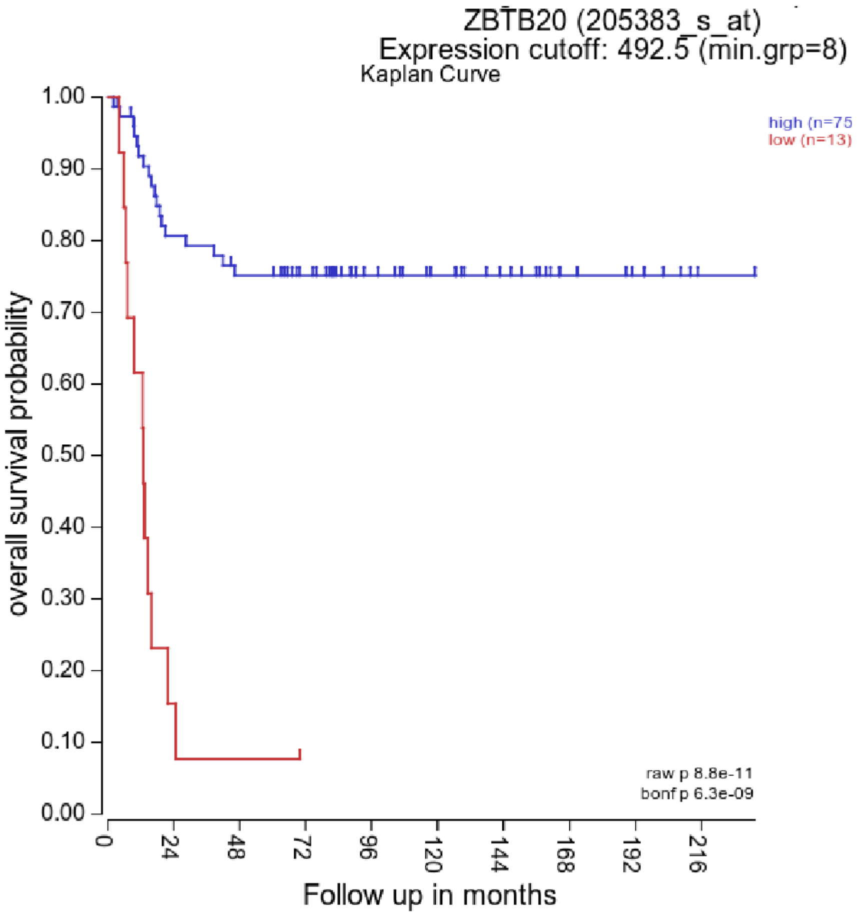


**Figure S9: Association between *ZBTB20* gene expression and survival.**
Kaplan-Meier curves with overall survival data of neuroblastoma samples from a publicly available dataset [51] separated into high and low expression of *ZBTB20* based on the threshold obtained with KaplanScan. Data and plots are retrieved from the R2 platform [66].

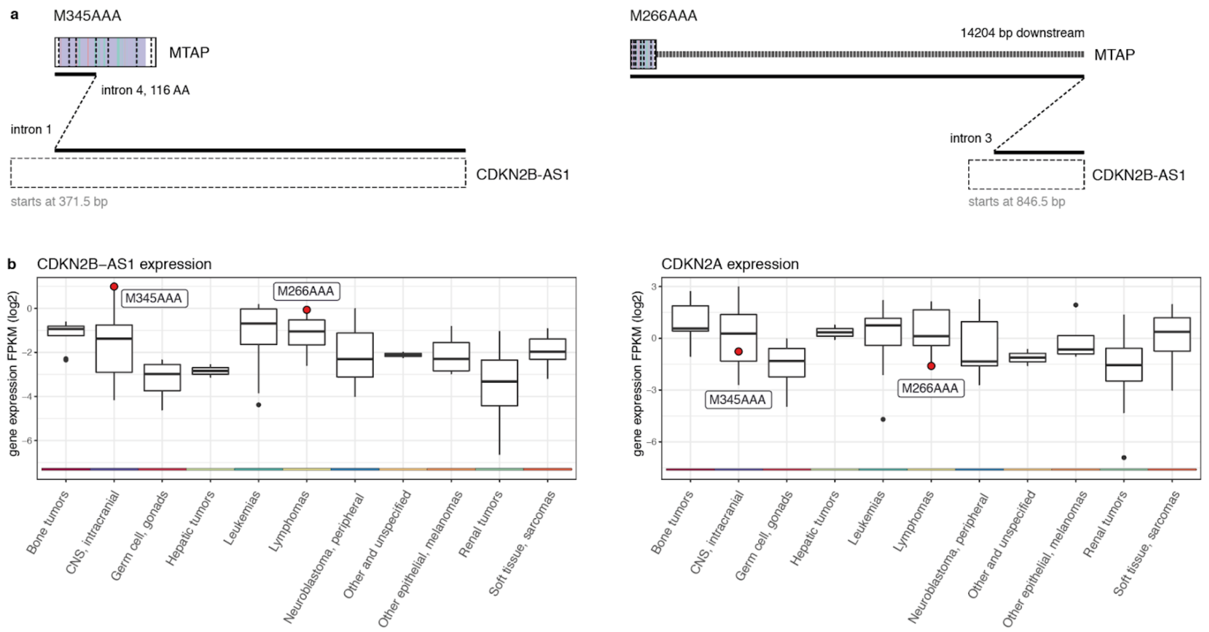


**Figure S10: *MTAP--CDKN2B-AS1* fusions and associated expression changes of *CDKNB-AS1* and *CDKN2A***

**a** Fusion schematics of *MTAP--CDKN2B-AS1* fusions generated using ProteinPaint from St. Jude Cloud [64, 65] from patients M345AAA (left) and M66AAA (right). **b** Gene expression levels (log2 FPKM) of *CDKNB-AS1* (left) and *CDKN2A* (right) for patient M345AAA and M66AAA carrying a *MTAP--CDKN2B-AS1* gene fusion (red circles) and all patients split according to their primary cancer type group (violin plots).
